# Precise calcium-to-spike inference using biophysical generative models

**DOI:** 10.1101/2024.12.31.630967

**Authors:** Gerard Joey Broussard, Giovanni Diana, Francisco J. Urra Quiroz, B. Semihcan Sermet, Nelson Rebola, Sanjeev Janarthanan, Laura A. Lynch, David M. Turner, David A. DiGregorio, Samuel S.-H. Wang

## Abstract

The intramolecular dynamics of fluorescent calcium indicators distort the relationship between calcium signals and action potentials (“spikes”), hampering efficient spike inference from calcium imaging. To address this problem, we characterized the calcium response kinetics of three widely used indicators, GCaMP6f, jGCaMP7f, and jGCaMP8f, using *in vitro* stopped-flow measurements and brain slice recordings. We identify previously unreported kinetic features, including use-dependent slowing of fluorescence decay, that introduce systematic errors in linear model-based inference methods. Using these observations, we developed a multistate model of GCaMP and used it to create biophysically-inspired Bayesian Sequential Monte Carlo and machine learning inference models trained on synthetic datasets. These methods outperform existing methods on spike timing accuracy and correlation benchmarks derived from diverse cell types. Our results show that using synthetic data derived from our biophysical model yields a decoder that outperforms even those trained on extensive experimental data. By separating indicator characterization from inference, our framework, **C**alcium **S**pike **P**rocessing using **I**ntegrated **K**inetic **E**stimation and **S**imulation (C-SPIKES), provides a generalizable strategy applicable to existing and future calcium indicators.

## Introduction

Genetically encoded calcium indicators (GECIs), including the widely used GCaMP family ^1–5^, enable large-scale recordings of neural population activity and are commonly used to generate a proxy variable for spike activity. However, the relationship between fluorescence signals and underlying action potentials is strongly distorted by the temporal filtering imposed by calcium dynamics and indicator kinetics. Spike-evoked calcium transients evolve on timescales that are slow relative to membrane voltage changes ^6^, and the subsequent calcium-to-fluorescence transformation introduces additional nonlinearities and delays ^7^. These combined processes limit the temporal precision of inferred neural activity and introduce systematic errors in spike reconstruction ^8–10^. In principle, accurate spike inference may benefit from models that explicitly account for the biophysical transformations that link spikes, calcium, and fluorescence.

A variety of supervised and unsupervised computational strategies have been developed to address the challenge of inferring spikes from fluorescence signals. Supervised approaches leverage data in which both fluorescence and spike times are known to train machine learning models to recognize statistical regularities that map observed signals to underlying spiking activity ^11,12^. Unsupervised approaches do not require labeled data, but instead rely on model-based assumptions about how spikes generate fluorescence to infer latent spike trains (e.g.,^13–19^. The performance of both categories improves when the relationship between spike occurrences and the resulting fluorescence responses is fundamentally linear ^16,20^.

In practice, this relationship often deviates from the linear ideal due to indicator saturation ^21^, calcium buffering ^22^, and supralinear indicator responses ^23^. Such nonlinearities degrade the accuracy of both simple unsupervised linear deconvolutions and supervised models that fail to generalize beyond their specific training regimes ^24,25^. Consequently, nonlinearities in the spike-to-fluorescence transformation have the potential to degrade the accuracy and reliability of inferred spike statistics across both method classes.

While recent jGCaMP8 sensors offer faster rise times and improved single-spike signal-to-noise ratios (SNR) ^5,26^, their inherent non-linearities remain unclear. Previous indicators exhibit supralinear responses to spikes that occur in rapid succession ^3,10,23,27–29^. Surprisingly, biophysical models incorporating this supralinearity have not outperformed simple linear approximations in spike decoding ^15,19^, suggesting the kinetic properties of even older sensors remain under-determined.

Here, we characterized the calcium-dependent kinetics of GECIs and integrated them into a biophysical framework for extracting spikes from fluorescence observables. We identified novel nonlinear properties in these responses, including a pronounced use-dependent slowing of jGCaMP8f during heightened activity. Using these insights, we developed a biophysical model of four GCaMP variants to power two new spike inference approaches: Biophys_SMC_, a sequential Monte Carlo (SMC) method for modeling latent spike-time sequences; and Biophys_ML_, a machine learning strategy trained on synthetic data generated using the SMC approach. Both methods outperformed state-of-the-art inference methods benchmarked on existing experimental datasets. Relative to other ML methods, BiophysML’s generative approach circumvents the need for simultaneous calcium fluorescence and ground truth spike recordings, enabling robust inference even without the need of the costly training data.

## Results

### Biophysical characterization of GCaMPs

To quantify reported kinetic and linearity improvements of jGCaMP8 variants ^5^, we expressed GCaMP6f, jGCaMP7f, or jGCaMP8f via viral vectors in cerebellar granule cells. We chose these cells because their bouton calcium transients summate linearly with spike number ^30^, allowing us to isolate indicator-specific nonlinearities. Using acute slices, we measured single-bouton fluorescence transients (**Figure 1A-C**) during extracellularly-evoked action potentials via high-speed (0.5-1 kHz) two-photon linescans. jGCaMP8f exhibits the fastest rise and decay times (**Figure 1C**), the largest fractional per-spike fluorescence change (ΔF/F) (**Figure 1D**; One-way ANOVA, F_(3,77)_ = 4.80, p = 0.004) and the highest signal-to-noise ratio (**Figure 1E**; One-way ANOVA, F_(2,70)_ = 28.99, p < 0.0001). Strikingly, jGCaMP8f’s rise time approached the near diffusion-limited kinetics of OGB-5N (DiGregorio et al., 1999) and was significantly faster than the other GCaMP probes (**Figure 1F**; One-way ANOVA, F_(3,77)_ = 28.99, p < 0.0001).

**Figure 1:**
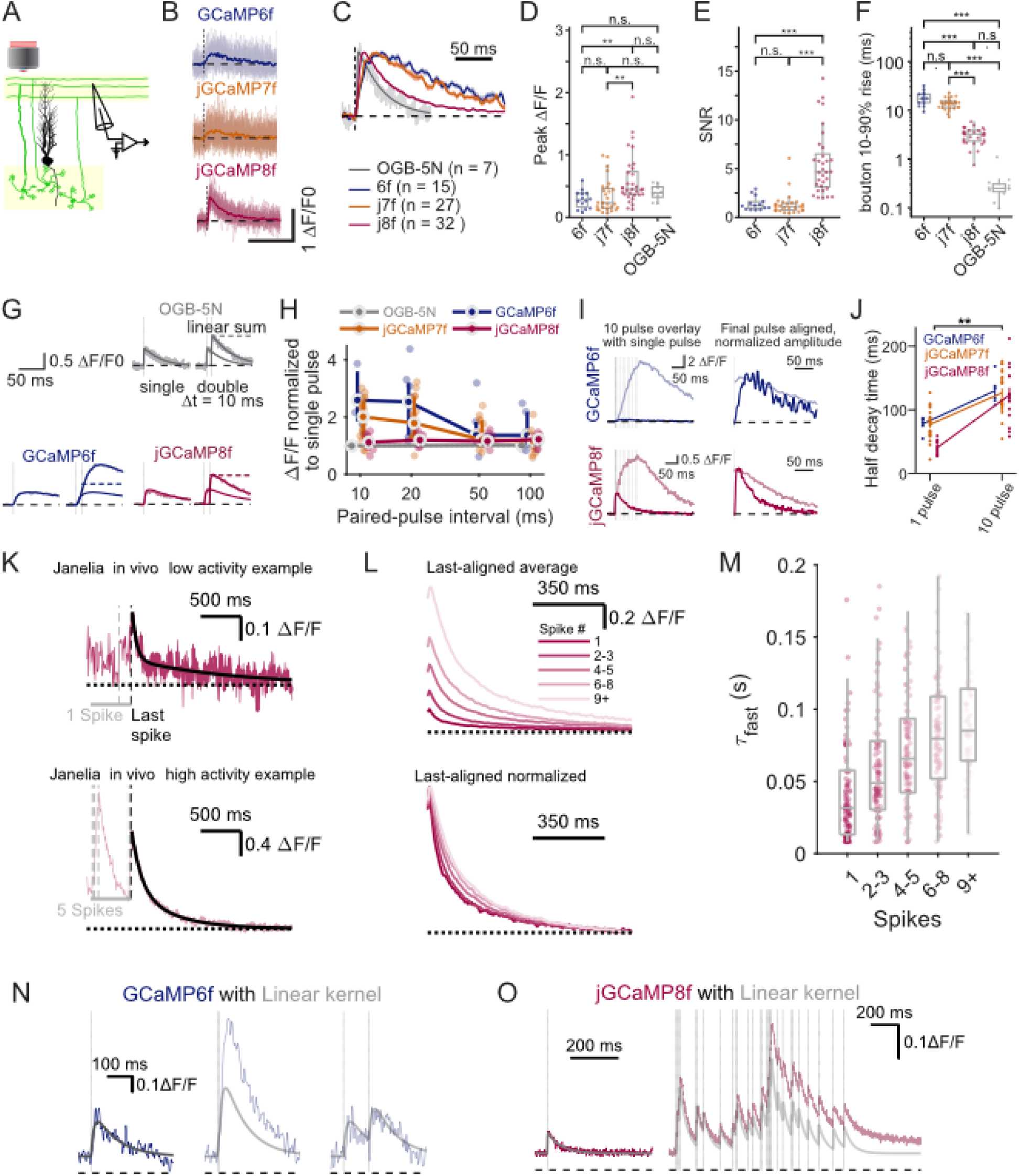
Nonlinear response characteristics of fast GCaMPs in a controlled, ex vivo setting. (**A**) Schematic of experimental approach showing sites of extracellular stimulation and imaged regions (highlighted in yellow). (**B**) Representative traces obtained across sensors for single-spike responses, with mean overlaid. (**C**) Traces from **B** averaged and normalized to compare the kinetic properties of GCaMPs with the low-affinity synthetic indicator, OGB-5N. (**D-F**) Peak response amplitude, signal-to-noise ratio, and 10-90% rise time for individual boutons. Each symbol represents the average across trials for a single bouton. Values reported as mean ± sem. (**D**) ΔF/F: jGCaMP8f, 0.59 ± 0.07, n = 32 boutons; GCaMP6f, 0.29 ± 0.04, n = 14 boutons; jGCaMP7f, 0.34 ± 0.05, n = 27; the small-molecule indicator Oregon Green BAPTA-5N, OGB-5N, 0.39 ± 0.04, n = 11. (**E**) SNR: jGCaMP8f, 5.6 ± 0.1; GCaMP6f, 1.4 ± 0.2; jGCaMP7f, 1.3 ± 0.2. (**F**) Rise times: jGCaMP8f, 3.1 ± 0.2 ms; GCaMP6f, 18.1 ± 1.6 ms; jGCaMP7f, 13.7 ± 0.7 ms; OGB-5N, 0.3 ± 0.1 ms; significant Tukey HSD post hoc comparisons: p = 1.1e-21, GCaMP6f vs. OGB5N; p = 5.4e-4, jGCaMP6f vs. jGCaMP7f; p = 3.9e-23, jGCaMP6f vs. jGCaMP8f; p = 8.7e-18, jGCaMP7f vs. OGB-5N; p = 1.5e-19, jGCaMP7f vs. jGCaMP8f. (**G**) averaged responses to one stimulus or two stimuli spaced 10 ms for the indicated sensors. (**H**) Relative amplitude of responses to a second stimulus spaced at various time intervals from the first stimulus. Fold increase: jGCaMP8f, 1.1 ± 0.1 at 10 ms and 1.2 ± 0.2 at 100 ms, n = 11 boutons; GCaMP6f, 3.1 ± 0.7 at 10 ms and 1.5 ± 0.3 at 100 ms fold increase, n = 7 boutons; jGCaMP7f, 1.9 ± 0.2 at 10 ms and 1.1 ± 0.1 at 100 ms, n = 14 boutons. Two-way mixed ANOVA, F(3,60) = 7.2, p = 3.3e-4 main effect of indicator, with significant one-way ANOVA at 10 ms, F(3,32) = 6.4, p = 0.002; significant Tukey HSD post hoc comparisons: p = 0.01, GCaMP6f vs. OGB-5N). Each point represents the average response for one bouton with the central emphasized point indicating distribution median and bar the IQR. Values reported as mean ± sem. (**I**) Responses to 1 stimulus and 10 stimuli at 100 Hz overlaid to the single stimulus response (left), and normalized and aligned to the last stimulus to demonstrate prolonged decay (right). (**J**) Half-decay times as a function of stimulus number. GCaMP6f, 78 ± 5 ms 1 pulse vs. 130 ± 9 ms 10 pulse, p = 0.029, n = 6 boutons; jGCaMP7f, 77 ± 5 ms 1 pulse vs. 126 ± 7 ms 10 pulse, p = 2.83e-6, n = 22 boutons; jGCaMP8f, 41 ± 3 ms 1 pulse vs. 123 ± 12 ms 10 pulse, p = 8.50e-10, n = 14 boutons; mean ± SEM. p-values calculated with Tukey’s HSD post hoc test. Error bars indicate standard error of the mean. (**K-M**) Demonstration of the elongation effect from **I-J** in vivo. (**K**) Examples of the fluorescence tail following low or high numbers of spikes. (**L**) Aligning of fluorescence responses by binned spike number in the 500 ms preceding the period shown. The bottom panel shows the alignment of normalized traces, with progressively higher activity demonstrating the in vivo elongation effect. (**M**) Boxplot of fit fast time constant (see Supplementary Figure 1 and Methods) of individual events binned by spike number in the count window. Mixed effects model included 1357 observations from n = 142 epochs. (**N-O**) Convolution of a single-stimulus response with the times of stimuli to demonstrate the effect of short-term amplitude non-linearity in GCaMP6f responses in **N** and cumulative signal arising from many stimuli for jGCaMP8f in **O**. Stars indicate statistical significance as: *p<0.05, **p<0.01, ***p<0.001.

Previous studies report a nonlinear relationship between spike number and GCaMP6 fluorescence^5,10,28^. Testing paired spikes across short intervals (10-100 ms), we found that GCaMP6f and 7f exhibit short-term supralinearity that declines to near-linear summation for intervals longer than 100 ms (**Figure 1G, H**; see figure legend for detailed Two-way mixed ANOVA and post-hoc statistics). In contrast, jGCaMP8f maintained near-linear response amplitudes across all intervals (**Figure 1G, H**; One-sample t-test, p = 0.32).

To examine the linearity of responses to bursts of activity, we measured bouton fluorescence in response to 1, 2, and 10 AP stimuli delivered at 100 Hz. GCaMP6f responses showed a strong supralinearity, in which a 10-pulse train evoked signals approximately 20-fold larger than a single pulse, consistent with positive cooperativity. In contrast, jGCaMP8f responses under the same comparison resulted in only a 3-fold larger response, consistent with indicator saturation (**Figure 1I, J**, bottom).

Unexpectedly, ten-spike bursts significantly slowed fluorescence decay across all tested GCaMPs compared to single spikes (**Figure 1I, J**; two-way mixed ANOVA, main effect of time F_(1,78)_ = 72.2, p < 0.0001). This use-dependent slowing was more pronounced in jGCaMP8f (∼3-fold vs. ∼1.5 in older variants) (**Figure 1J**). Fitting the jGCaMP8f decay required a two-component exponential even for single spikes, with the fast component in particular lengthening as spike count increased (**Supplementary Figure 1A-D**), independent of saturation (**Supplementary Figure 1E-G**). Ground truth *in vivo* recordings ^5^ confirmed this progressive decay slowing for jGCaMP8f (**Figure 1K, L**), specifically affecting the faster time constant *τ*_1_ of the decay process (**Figure 1M**, Linear mixed effects model, fixed effect of *τ*_1_; F_(1,240.8)_ = 105.1, *β* = 0.10, p<0.0001). Thus, jGCaMP8f exhibits activity-dependent slowing in the fluorescence decay.

To assess the impact of nonlinearities during *in vivo*-like activity bursts, we compared measured spike responses against a linear kernel summation model (convolving a single-spike kernel with the stimulus spike train; see **Methods**). For paired spikes, the GCaMP6f model underestimated responses at 10 ms intervals, improving only as intervals lengthened (**Figure 1N**). As our observation of elongated decay suggested, exposing jGCaMP8f boutons to a 10 Hz Poisson train caused the kernel estimate to progressively diverge from the observed data during periods of high spiking activity (**Figure 1O**). These findings show that across GCaMP sensors, high activity regimes cause sensor response properties to diverge from the linear ideal.

**Supplementary Figure 1:**
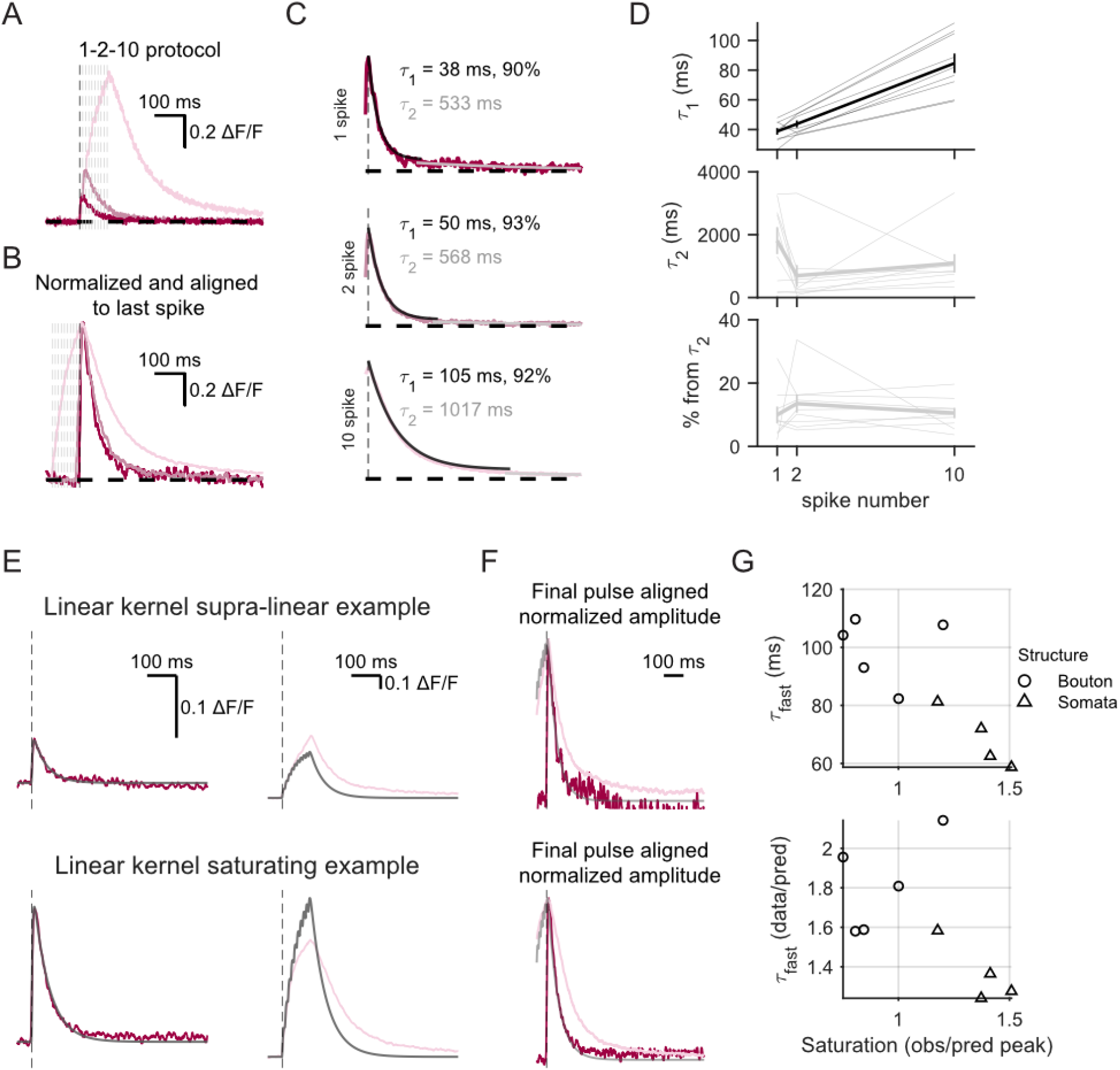
jGCaMP8f response decay is captured by two kinetic components, while its slowing cannot be explained by response saturation. (**A**) Representative single-bouton responses to 1, 2, and 10 stimuli overlaid and aligned by first stimulus time. (**B**) Responses normalized and aligned to last spike. (**C**) Example fits to dual sums of exponentials showing fast and slow components of decay. (**D**) Dependence of fast (top) and slow (middle) time constants on the number of stimuli, and relative contribution of the slow component (τ_2_) to the overall time course (bottom). Individual bouton values shown as low linewidth traces with group means shown as thicker line and standard error of the mean as error bars, n = 6 boutons and 3 granule cell somata. (**E**) Sample responses from Purkinje granule cell soma (top) and bouton (bottom) with a linear fit to their respective single pulse response (left) and the convolution of the unitary response overlaid to the fluorescence response for each structure showing that GCaMP in the soma was operating in a non-saturating response regime while the bouton was beyond the linear regime. (**F**) Normalized alignment to the last pulse response in sub- (top) and supra-saturating regimes that demonstrate elongation of fluorescence response in the 10 pulse response in each case. (**G**) Fluorescence response elongation increases with cumulative calcium load as indexed by degree of supra-versus linear response (top) while the elongation is greater than predicted by the linear response (bottom).

**Supplementary Figure 2:**
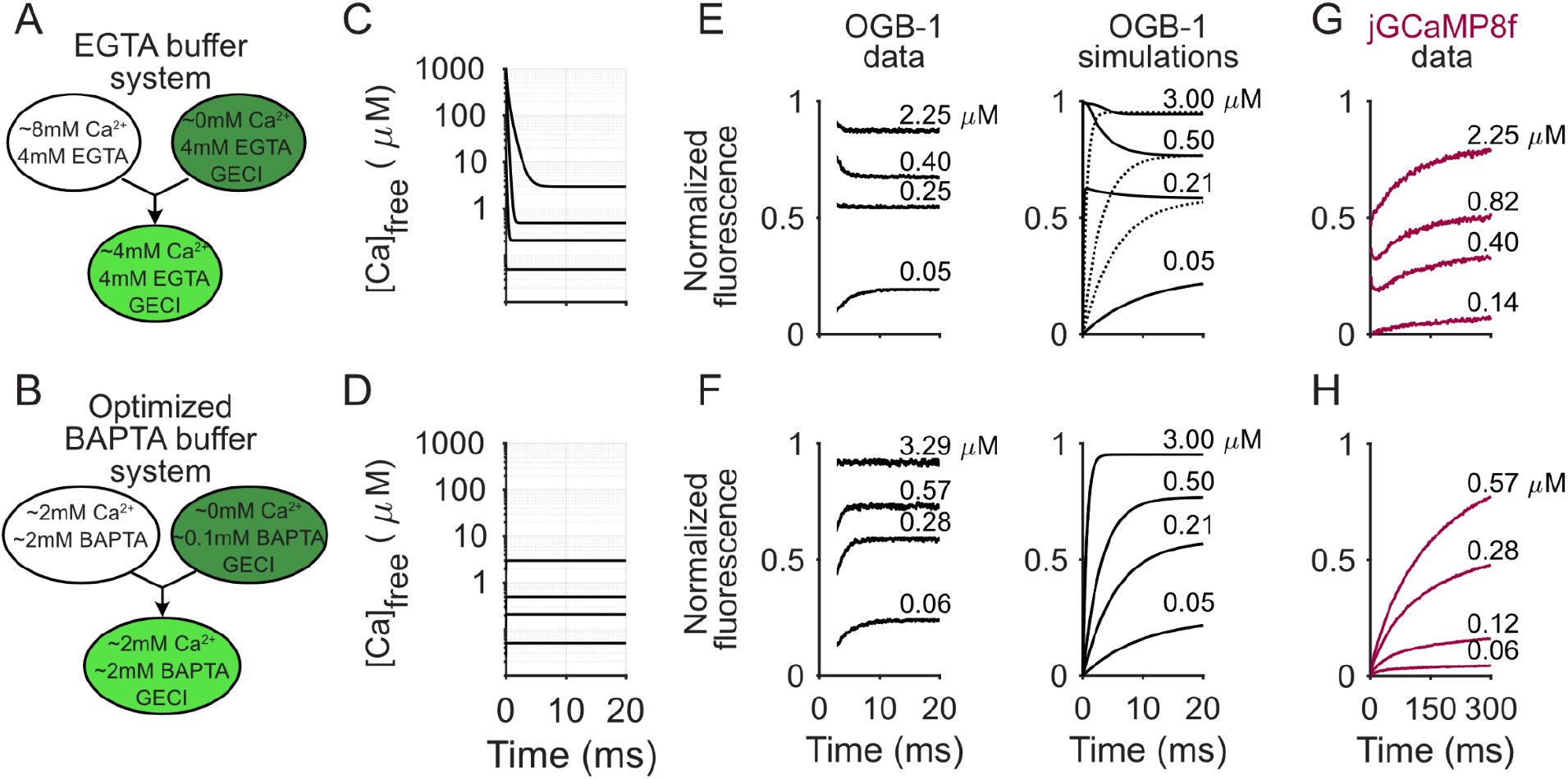
The impact of calcium clamp design on observed stopped-flow dynamics. (**A-B**) Schematic of stopped-flow buffer systems using (**A**) a slow buffer (EGTA) and containing unbuffered calcium in the calcium-containing bolus (**A**) and (**B**) a system with a fast buffer (BAPTA) and fully buffered calcium in the calcium-containing bolus. (**C**) Simulations of the EGTA-based system presented in (**A**) show uncontrolled free calcium concentrations in the first few milliseconds of large calcium steps, but (**D**) BAPTA-based experiments do not. (**E**) Measurements of OGB-1 responses (left) reflect uncontrolled responses that are matched by simulations of the buffer system (right). (**F**) OGB-1 data and simulations in BAPTA-based buffers. (**G**) Artifactual fast on-response to jGCaMP8f in EGTA-based buffer system. (**H**) True jGCaMP8f response in BAPTA-based buffer system.

**Supplementary Figure 3:**
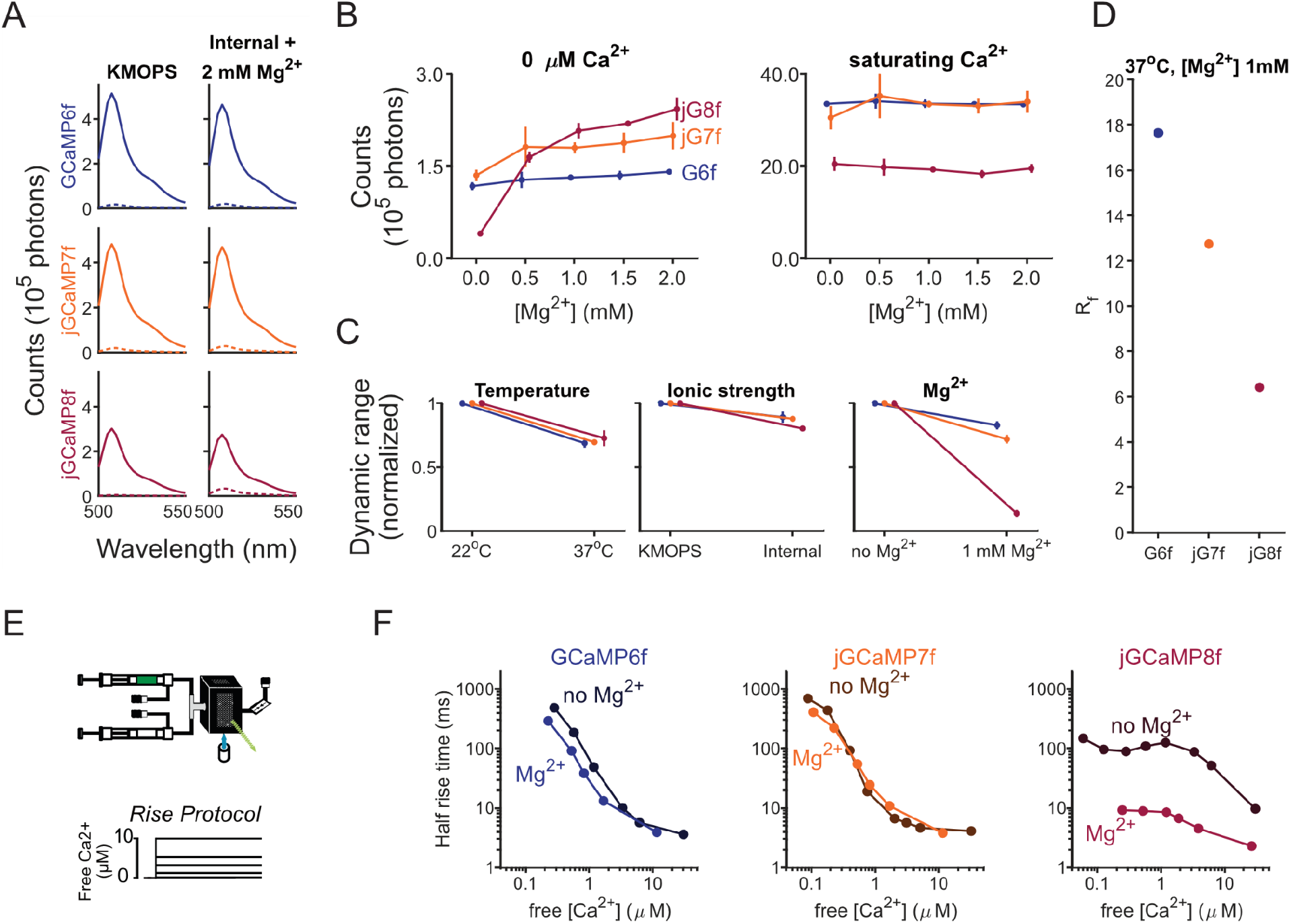
Magnesium affects the properties of jGCaMP8f much more than older GCaMPs. (**A**) Emission spectra of GCaMP variants in standard protein purification buffer (KMOPS, left) and under intracellular conditions that include magnesium (2 mM Mg^2+^). (**B**) Fluorimeter counts at 490 nm under calcium-free (left) or saturating calcium (right) conditions. jGCaMP8f, but not other variants, showed strong magnesium-dependence of fluorescence emission under low-calcium conditions. No indicator showed this effect at saturating calcium. (**C**) Quantification of the weak dependence of GECI dynamic range R_f_ (saturating calcium divided by calcium-free fluorescence output) on temperature (left) or ionic strength (center) compared to the effect seen with magnesium (right). (**D**) Dynamic range of GCaMP variants at mammalian near-physiological conditions. (**E**) Schematic of stopped-flow experiment to measure responses to rising steps of calcium concentration. (**F**) Half-rise times to calcium steps were accelerated most by magnesium for jGCaMP8f, less so for GCaMP6f, and not at all for jGCaMP7f. Error bars indicate standard error of the mean.

To investigate the mechanistic basis of GCaMP’s use-dependent nonlinearities, we measured purified protein fluorescence responses to calcium concentration steps using stopped-flow fluorimetry (**Figure 2A-D**). Optimizing the buffer to match intracellular ionic composition (including magnesium) was necessary to reproduce neuronal kinetics and dynamic range (**Supplementary Figures 2 and 3**). Across 0-10 μM calcium steps, jGCaMP8f exhibits faster rise and decay kinetics than 6f and 7f (**Figure 2C, D**). For calcium increases corresponding to a small number of spikes (shaded region, **Figure 2E**), stopped-flow rise times correlated tightly with spike-evoked calcium transient rise times across variants (**Figure 2F**; linear regression, R^2^ = 0.99, p = 0.017). Additionally, analysis of the early phase of stopped-flow responses showed jGCaMP8f has the lowest cooperativity (Hill coefficient 1.35 vs. > 2; **Figure 2H**). These observations align with the linear paired-pulse responses of jGCaMP8f and the supralinear responses of 6f and 7f (**Figure 2G-I**; linear regression, R^2^ = 0.99, p = 0.001), demonstrating that our stopped-flow measurements accurately capture physiological spike-evoked kinetic features.

**Figure 2:**
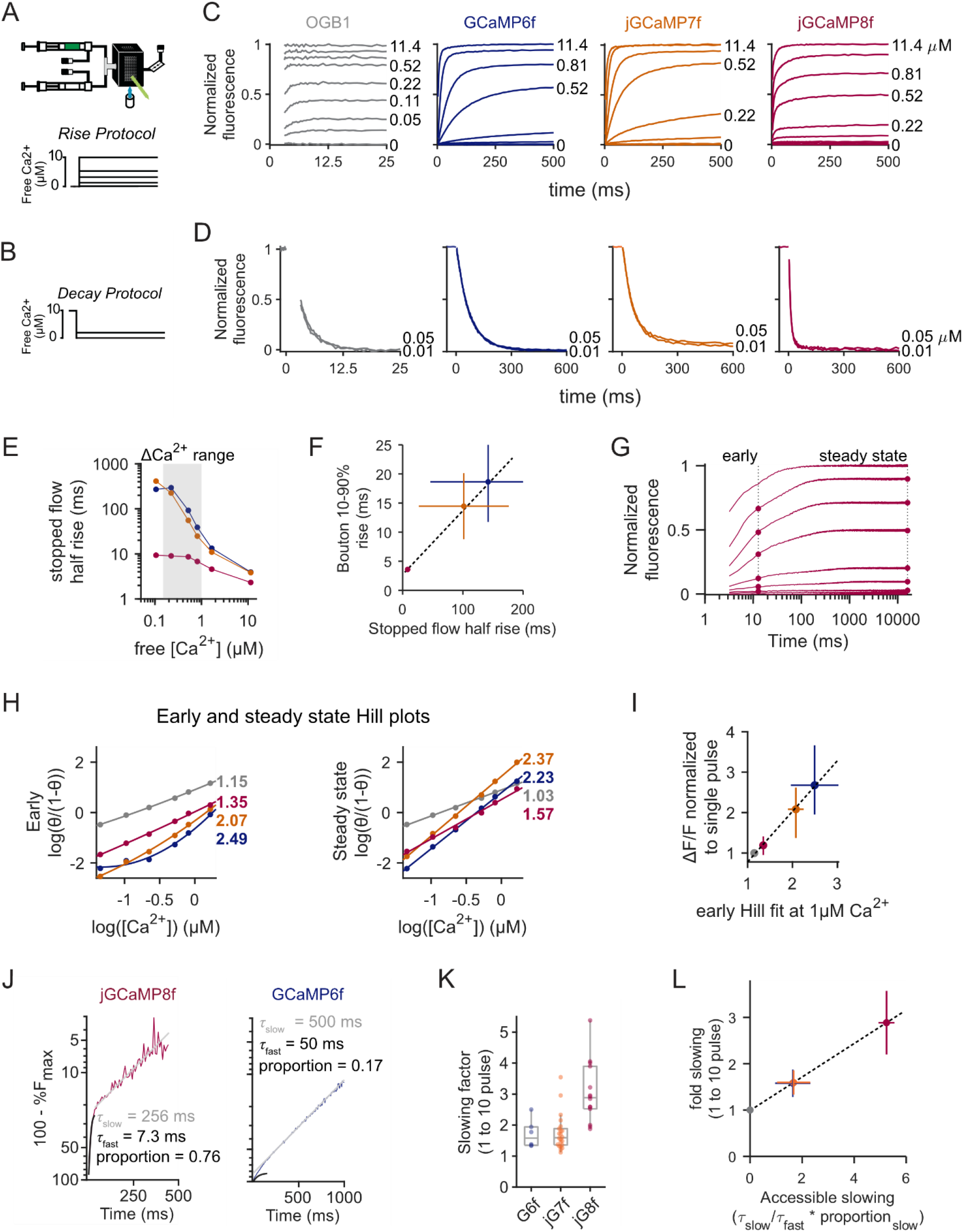
Stopped-flow fluorimetry of GCaMP dynamic responses. (**A**) Schematic of stopped-flow experiment and rising steps of calcium concentration. (**B**) Step-down protocols for decreasing calcium. (**C**) Responses to steps up from 0 μM free calcium. (**D**) Responses to steps down from 10 μM free calcium. (**E**) Half-rise times for GCaMP6f, jGCaMP7f, and jGCaMP8f as a function of calcium step size. Shaded gray region indicates the physiological range of intracellular signals. (**F**) Comparison of brain slice presynaptic bouton signals with stopped-flow rise times for the three indicator variants (**G**) Rising phase responses on logarithmic time scales. Symbols indicate the time points plotted in (**H**). (**H**) Hill plots of rising and steady-state responses to steps in calcium. Colored numbers indicate Hill coefficients. (**I**) Comparison of brain slice amplitude nonlinearity at 10 ms inter-stimulus response with respect to Hill coefficients. (**J**) Representative stopped-flow traces with y-axis values presented on a log scale for jGCaMP8f (left) and GCaMP6f (right). Sums of exponentials for each indicator demonstrate the biphasic response and the greater contrast in speed of kinetic components seen in jGCaMP8f. (**K**) Quantification of the factor of slowing seen in GCaMP fluorescence decay in response to 1 and 10 stimuli in brain slices. Significant Tukey HSD post hoc comparisons include GCaMP6f vs. jGCaMP8f, p = 0.0015; jGCaMP7f vs. jGCaMP8f, p = 1.7e-5. (**L**) Comparison of brain slice slowing to accessible slowing (see main text and methods). Error bars in all panels represent the interquartile range of the distribution.

Consistent with prior studies showing fast and slow pathways for calcium-mediated fluorescence increases ^7,31^, calcium step responses across GCaMP sensors were distinctly biphasic. For GCaMP6f and jGCaMP7f, fluorescence changes occurred through a slow component that accelerated with larger calcium steps. In contrast, jGCaMP8f’s rising fluorescence developed largely through a fast component which showed a notably shallow dependence on calcium concentration (**Figure 2J, Supplementary Figure 4A-C**). Responses to calcium down-steps were similarly biphasic (**Supplementary Figure 4D-G**). The observed fast rise times at low calcium concentrations may explain jGCaMP8f’s faster *in vivo* rise time at the lower peak calcium concentrations typical of single spikes (**Figure 2J, Supplementary Figure 4**).

The biphasic rising and decay kinetics suggest calcium-driven entry into at least two distinct fluorescence states. We compared these fast and slow time constants against the use-dependent slowing observed in our *ex vivo* slice recordings. We found a linear relationship between our *ex vivo* data and “accessible slowing”, a metric representing the tradeoff between speed of entry into theoretical fast versus slow states and the stability of the slow state (**Figure 2K**; One-way ANOVA, F_(2,39)_ = 15.2, p < 0.0001, **2L**; linear regression, R^2^ = 1.00, p < 0.0001). These results are consistent with a model in which large calcium increases push GCaMP variants into a stable fluorescent state from which they are slow to exit.

**Supplementary Figure 4:**
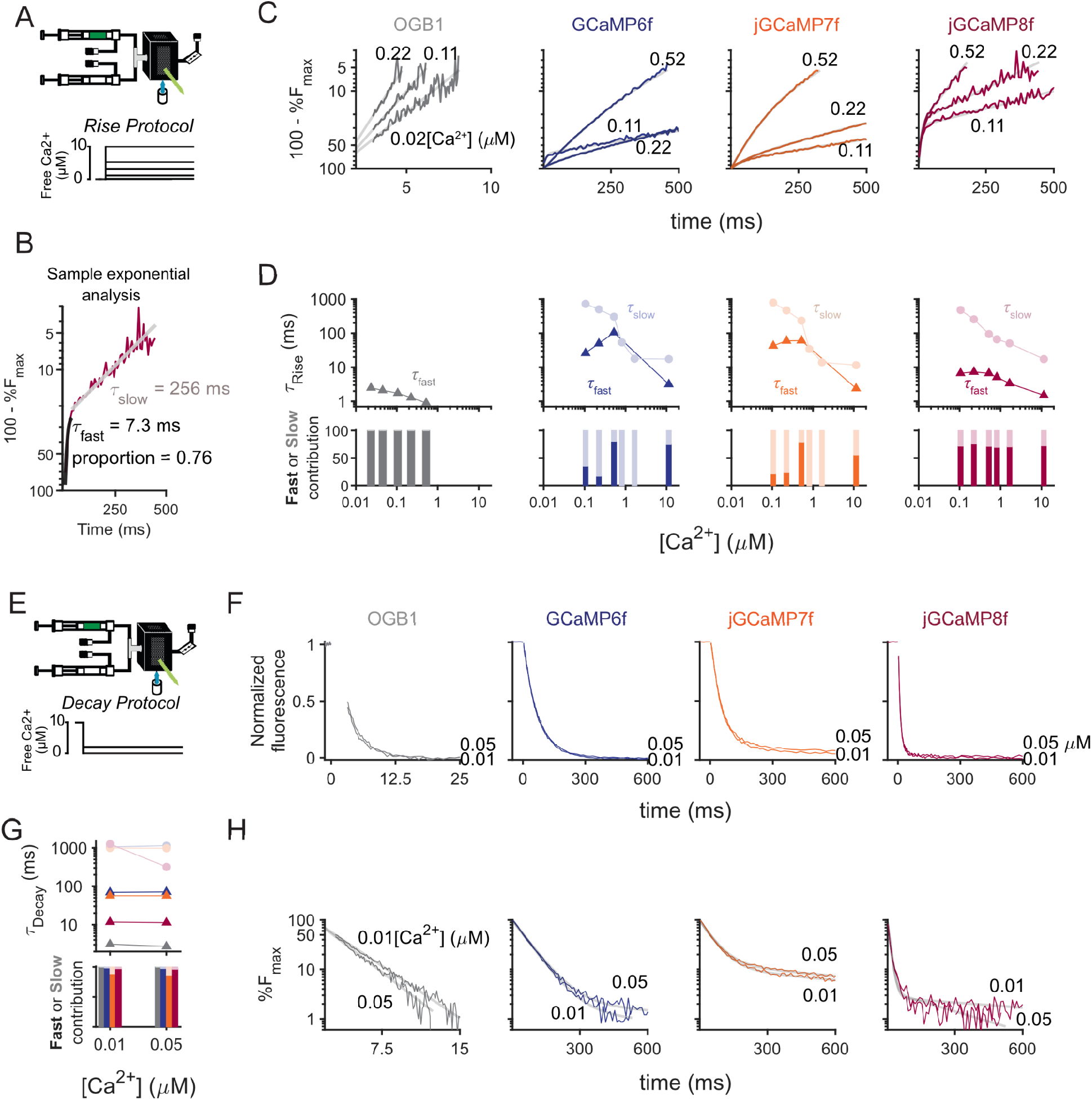
Biphasic responses of GCaMPs to increasing calcium steps. (**A**) Schematic of stopped-flow experiment and rising calcium-step protocol. (**B**) Example analysis of fit to a sum of two exponentials. (**C**) Monoexponential rising kinetics for Oregon Green BAPTA-1 (OGB-1) and biexponential kinetics for GCaMP variants. (**D**) Dependence of fast and slow components on amplitude of calcium step. Upper plots show rate constants represented in the response at the indicated step. In the lower plots, vertical bars show relative amplitude of fast (dark) and slow (light) components. (**E**) Schematic of decreasing-calcium step protocol. (**F**) Fluorescence responses to decreasing calcium steps. (**G**) Breakdown of kinetic components contributing to the decay of fluorescence as in (**D**). Due to the low number of steps considered, analysis for all variants is included in this single plot. (**H**) Monoexponential decay for OGB-1 and more complex responses for GCaMP variants.

**Supplementary Figure 5:**
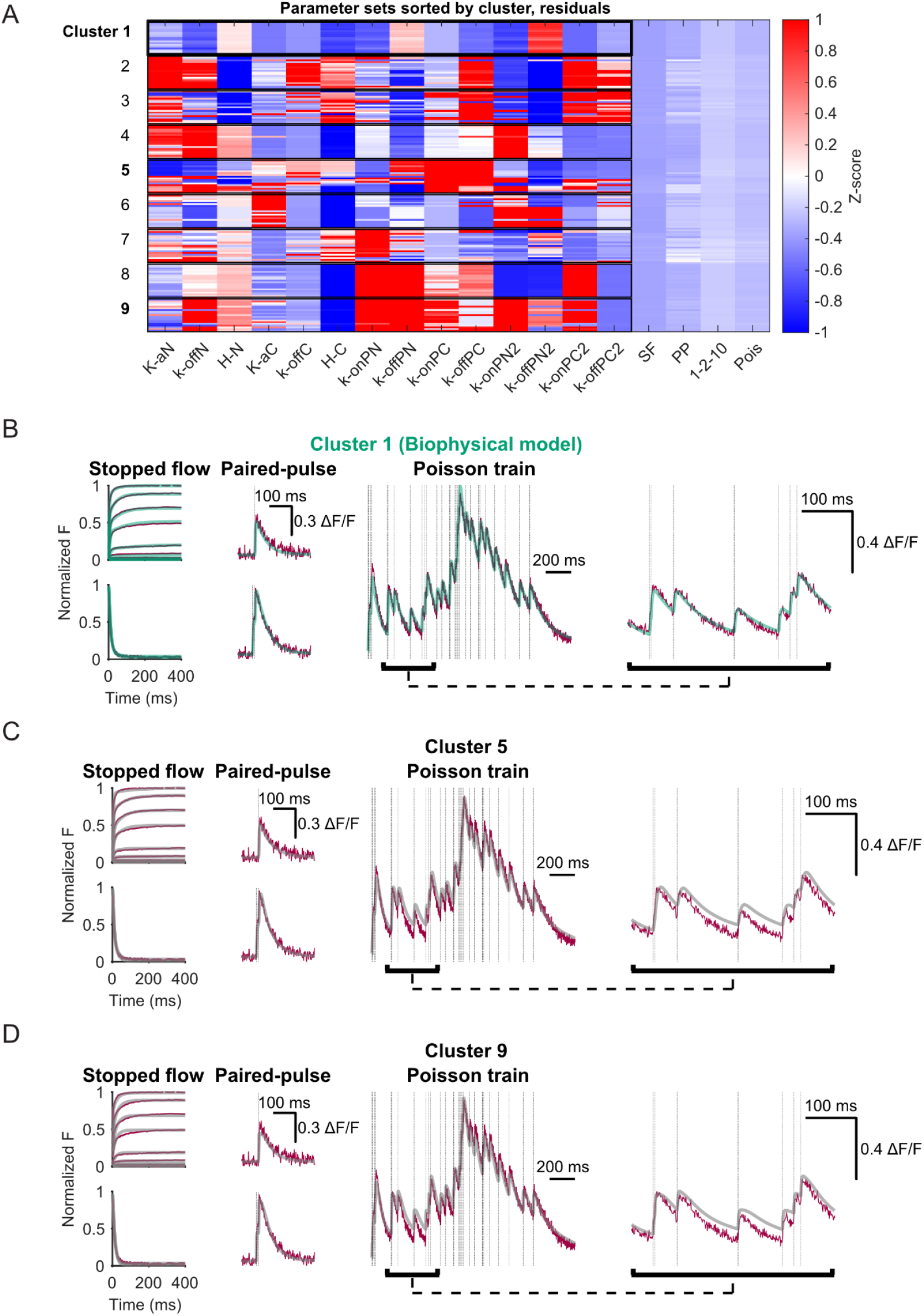
Selection of jGCaMP8f biophysical parameters using in vitro and ex vivo datasets. (**A**) Distributions of parameters on fits from randomized seeds with GCaMP model parameters at figure left and residuals from best fits to stopped-flow and ex vivo experiments (paired pulse, 1-2-10 pulses, and Poisson train) based on optimizing cell parameters. GCaMP parameter sets were sorted by hierarchical clustering and then by lowest summed residuals within each cluster. This procedure allowed qualitative assessment of parameters that lay in different local minima of the parameter space. (**B**) Fitted data examples for cluster 1, which was used for all further modeling. (**C**) Fitted data for cluster 5. (**D**) Fitted data for cluster 9. In paired-pulse and Poisson train data, vertical lines indicate the times of action potentials. Expanded timescale of Poisson responses at right. Neither cluster 5 or 9 captured the differential slowing across the Poisson responses.

### Development of a biophysical model of GCaMP

To quantify fast and slow fluorescence transitions and observed cooperativity, we developed a biophysical GCaMP model incorporating parallel fast and slow kinetic pathways. This design reflects calmodulin’s underlying calcium binding properties, where the N-lobe (fast) and C-lobe (slow) semi-independently bind calcium^32,33^ and subsequently interact with a target peptide to increase fluorescence^7,34–36^. We reasoned this structure could phenomenologically capture both *in vitro* and *ex vivo* observations (**Figure 3A**).

**Figure 3:**
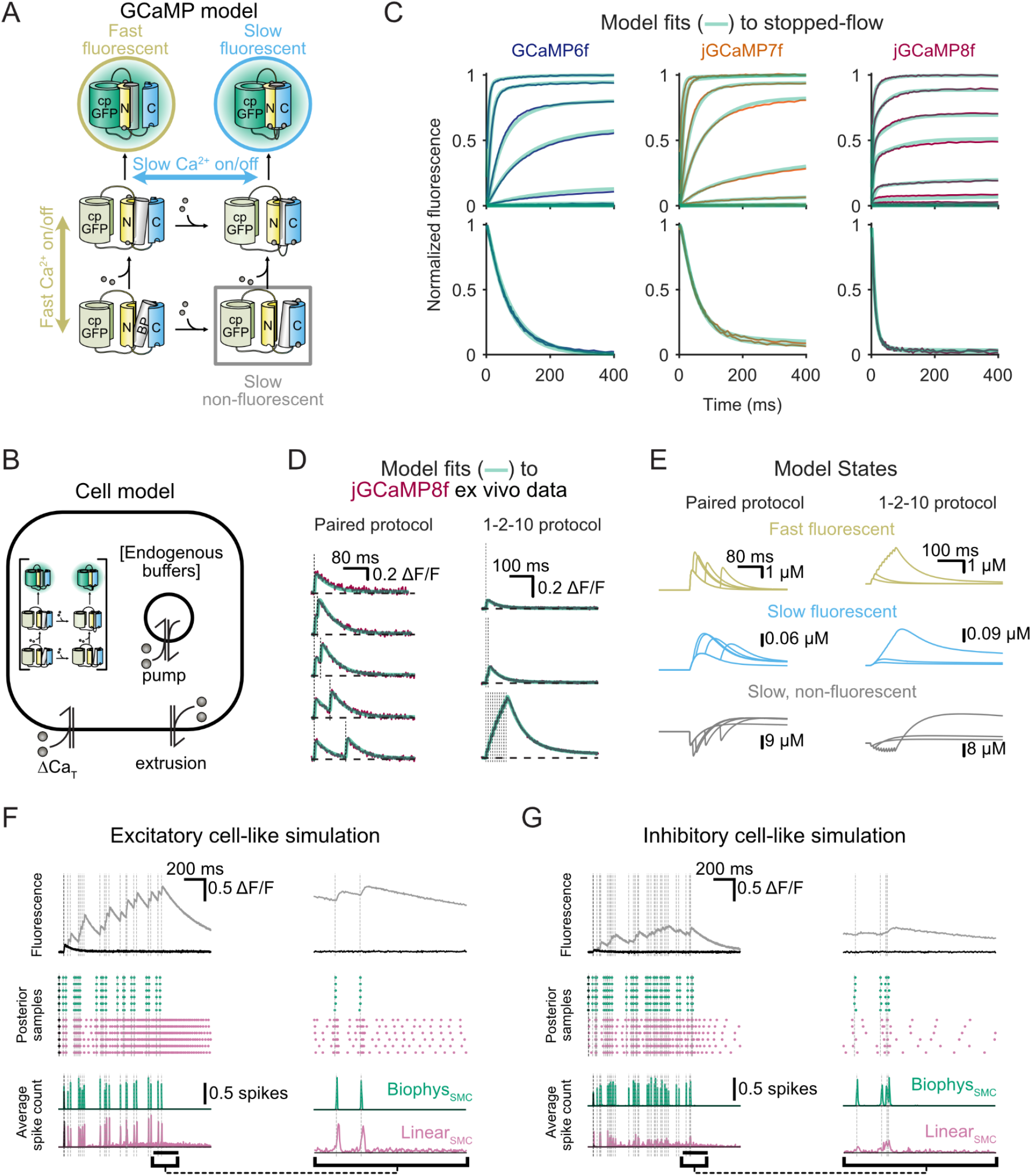
A biophysical model for GCaMP response. (**A**) Schematic GCaMP model depicting probe components, including circularly permuted green fluorescent protein (cp-GFP), the N- and C-lobes of the calcium binding-calmodulin, and the binding peptide (BP). Black arrows indicate key pathways to fluorescence with states highlighted for their role in model function. (**B**) Simplified schematic diagram of physiological response of a cellular structure containing endogenous calcium buffer, GCaMP, and intracellular and plasma membrane calcium extrusion (pump) mechanisms (see **Methods**). (**C**) Fits of the GCaMP model to the three GCaMP variants with different parameters for each variant. (**D**) Fits of the cell model to physiological data with fixed GCaMP model parameters. Vertical lines indicate action potential (spike) times. (**E**) Occupancy of the fast-fluorescent, slow-fluorescent, and slow non-fluorescent states of GCaMP indicating a mechanism for use-dependent slowing of fluorescence decay, becoming most apparent after 10 action potentials. (**F**) Predictions on simulated ground-truth data for an excitatory neuron. The data consists of a single spike (black) or a 10 Hz Poisson train of spikes (gray) (top panel). Lower panels show the average posterior spike probability (solid lines) and spikes times from a single trajectory (triangles) derived by our SMC method employing either the biophysical model (Biophys_SMC_, green) or a linear model (Linear_SMC_, pink) as its generative engine. Vertical lines indicate action potential (spike) times. (**G**) As in (**F**) with simulated ground truth for an inhibitory cell (30 Hz Poisson spike rate, calcium increase per spike reduced by 2, dynamic range reduced by a factor of 2).

We estimated GCaMP model kinetic rate constants by fitting *in vitro* (stopped-flow) data, constrained by *ex vivo* spiking responses from cerebellar granule cells (GCs), which have well-characterized calcium handling^37^. Our cell model built on the molecular model by incorporating endogenous buffering, extrusion, and intracellular store exchange (**Figure 3B**). For each variant, we generated candidate GCaMP models according to their stopped-flow fits, fixed GCaMP parameters, and fit the cell model to our GC dataset. The final, fixed parameters successfully accounted for dynamics across both datasets (**Supplementary Figure 5; Supplementary Figure 6A, B**), allowing us to reproduce variant-specific dynamics using a unified model structure (**Figure 3C, D; Supplementary Figure 6C-F**).

A close examination of the occupancy of each state of the GCaMP model (**Figure 3A; Supplementary Figure 6A**) revealed how parameterization dictates distinct operational modes. Categorizing fluorescent states by their dependence on C-lobe-like binding (slow) versus independence (fast), we found that only jGCaMP8f exhibited stable occupancy of fast fluorescent states across both *in vitro* and *ex vivo* fits (yellow traces, **Supplementary Figure 6C-F**). During decay, jGCaMP8f’s response slowing after spike bursts was driven by increased occupancy of slow fluorescent and non-fluorescent states (**Figure 3E**, blue and gray).

**Supplementary Figure 6:**
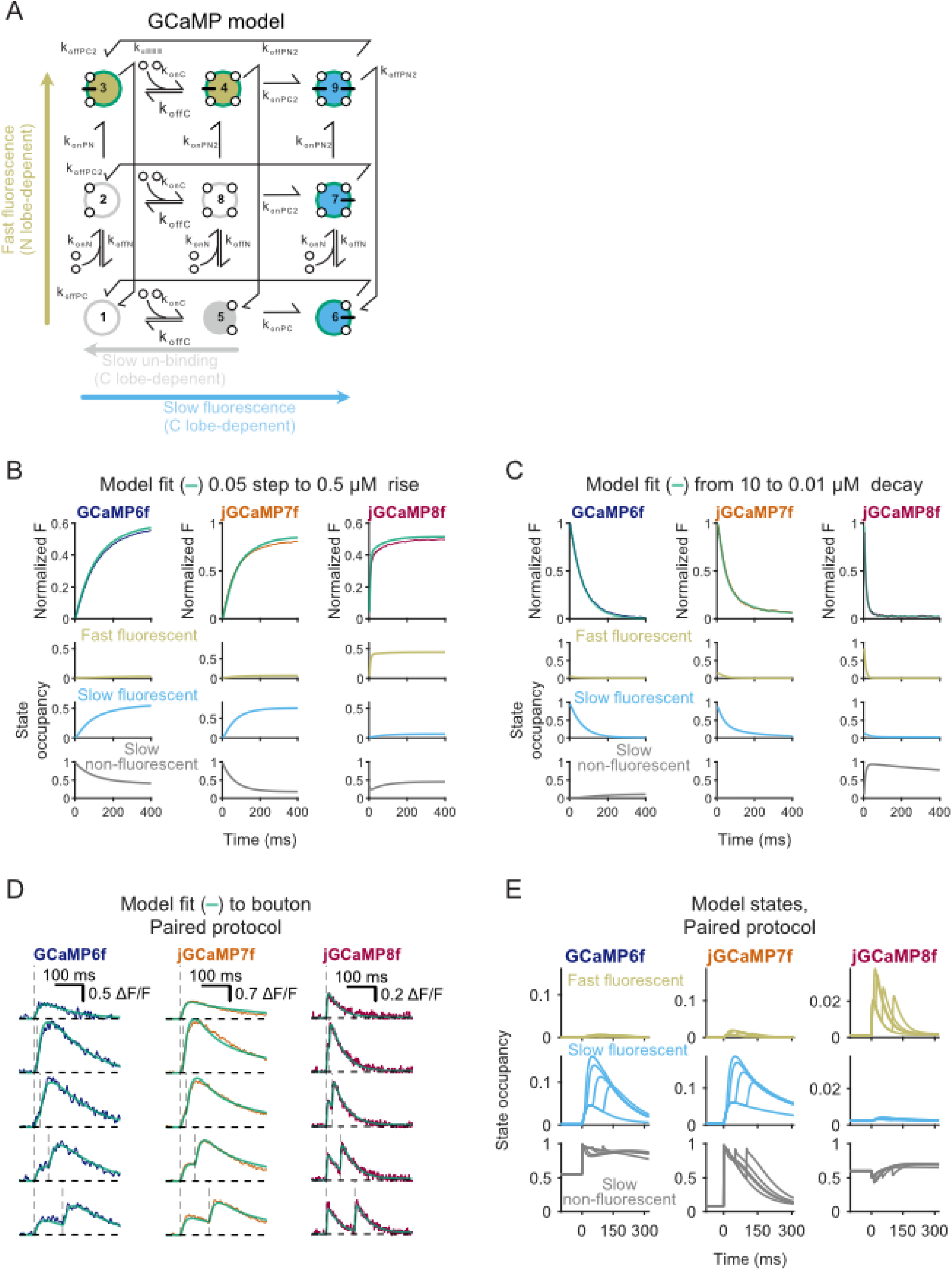
A full biophysical state analysis of GCaMP response. (**A**) Full biophysical state model of GCaMP response. Specific states referred to as “fast fluorescent”, “slow fluorescent”, and “slow, non-fluorescent” have centers color-coded as in model state traces below. (**B**) Model fits to 0.5 μM increasing step of calcium (top), with model state broken down by occupancy. Note that jGCaMP8f is distinguished by its near-exclusive use of the fast fluorescent states. (**C**) As in (**B**) for decreasing step. jGCaMP8f is distinguished by fast, persistent entry into the slow, non-fluorescent state. (**D**) Model fits to fluorescence responses in presynaptic boutons of granule cells. (**E**) State occupancy for the fluorescence responses shown in (**D**). Fast responses arise in part from high occupancy of a slow, non-fluorescent state at basal calcium for jGCaMP8f which efficiently transitions to fluorescence on elevation of calcium.

**Supplementary Figure 7:**
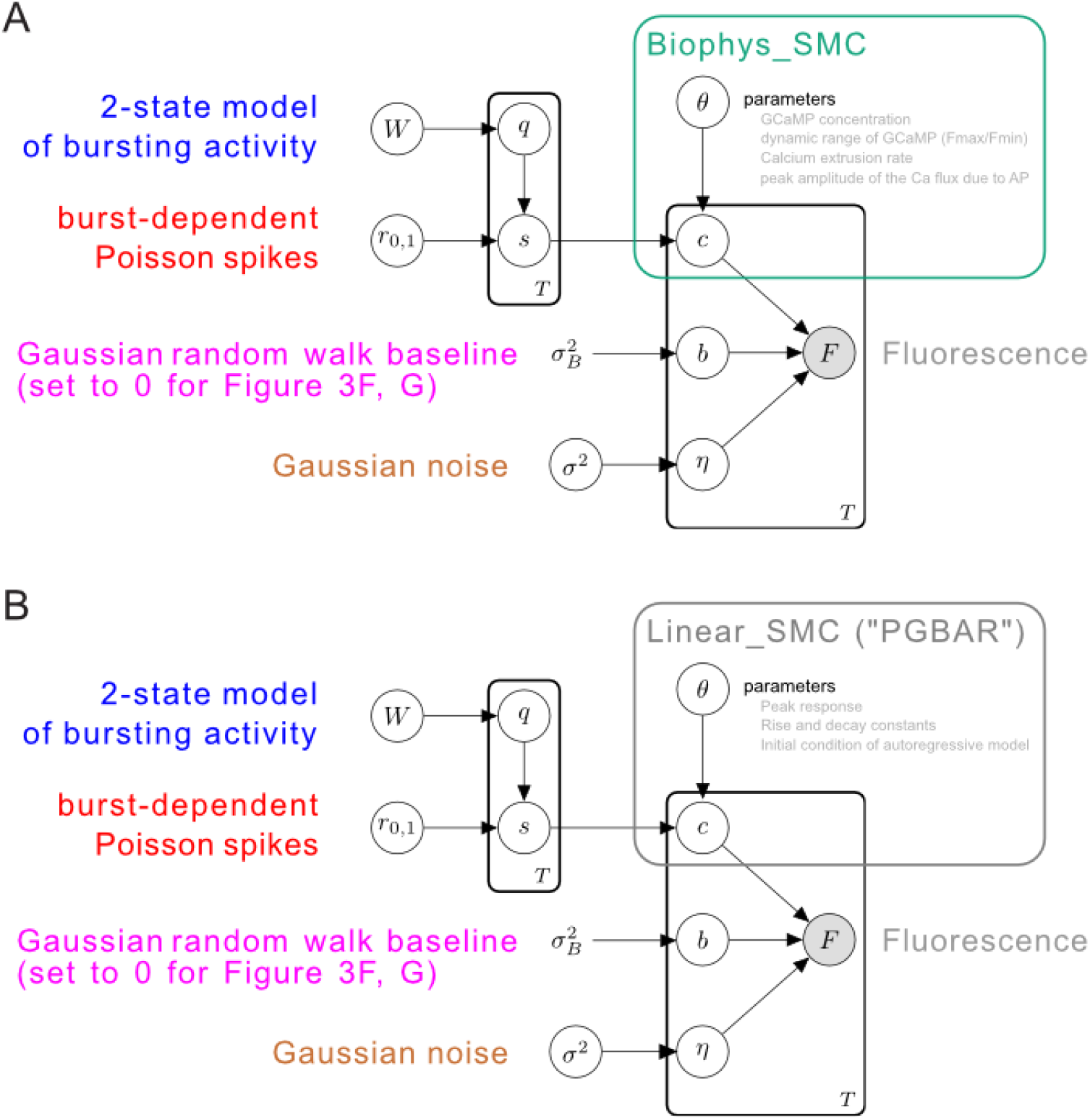
Generative cell-based model of fluorescence time series. (**A**) Graphical representation of the generative SMC model described in the main text. White circles denote unknown variables, gray circles denote measurements and the bare variable σ^2^_B_ is a fixed prior hyperparameter controlling the Brownian motion of the fluorescence baseline. This value was held fixed at 0 for the demonstration in **Figure 3F, G**. Rectangular outlines denote groups of variables. Note that GCaMP model parameters are included, but other than concentration are fixed for the SMC approach. (**B**) The key change over Linear_SMC/PGBAR (Diana et al., 2026) is the replacement of the autoregressor model with a biophysically-inspired parametric model (green rectangle in BiophysicalSMC, gray in Linear_SMC/PGBAR).

**Supplementary Figure 8:**
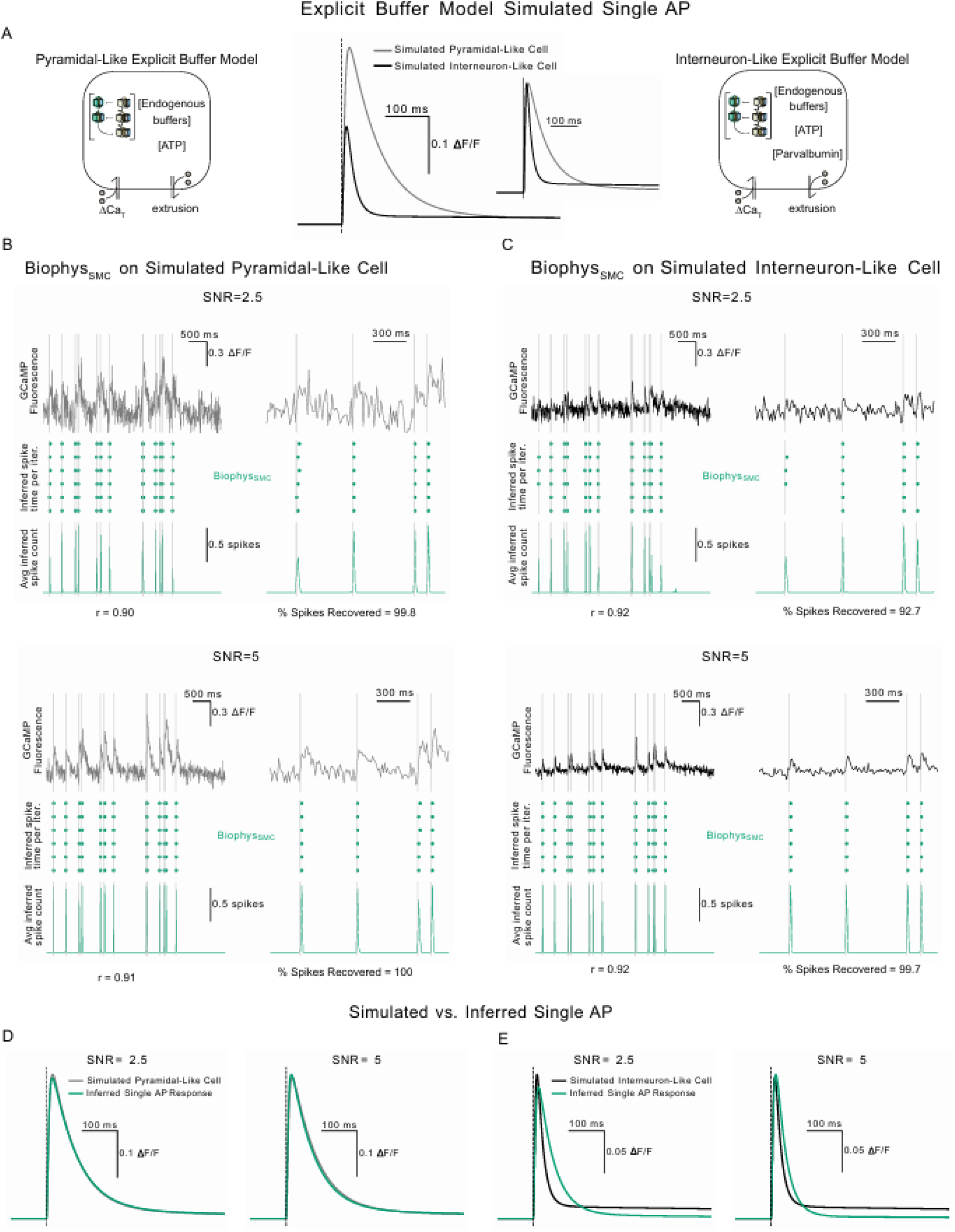
Biophys_SMC_ accurately predicts spikes in simulations of cell types with a wide variety of calcium handling properties. (**A**) Simplified schematic diagrams of explicit buffer models used to generate GCaMP responses with (interneuron-like) and without parvalbumin (pyramidal-like) (see Methods). Example simulated single action potential fluorescence responses for the two cell types are shown, along with normalized ΔF/F traces. (**B**) Spike inference using Biophys_SMC on simulated pyramidal-like GCaMP fluorescence traces driven by a 10 Hz Poisson train of spikes at a duty cycle of 50% and signal-to-noise ratios (SNRs) of 2.5 and 5. The top row for a given SNR shows simulated fluorescence traces generated by the explicit buffer model. Middle row shows posterior spike samples from individual iterations. Bottom panels show the average inferred spike count. The left column shows 5 seconds out of the full 60 second simulation and the right column shows the first second of that window. Dashed vertical lines indicate the ground-truth spike times used to generate the fluorescence traces. Correlation between inferred and ground-truth spike counts and the percentage of spikes recovered are indicated. (**C**) Same analysis as in (**B**) for simulated interneuron-like cells driven by the same Poisson spike train. (**D**) Comparison between simulated and inferred single action potential fluorescence responses for pyramidal-like cells at SNRs of 2.5 and 5. The inferred kernel was generated using the inferred cell model parameters from Biophys_SMC. (**E**) Same comparison as in (**D**) for simulated interneuron-like cells.

### Biophysical models for high-fidelity spike inference

We next leveraged our biophysical model of jGCaMP8f as a generative model core for two new spike inference methods: Biophys_SMC_ (a sequential Monte Carlo approach) and Biophys_ML_ (a machine learning strategy trained on synthetic SMC data). Together, these comprise the **C-SPIKES** (**C**alcium **S**pike **P**rocessing using **I**ntegrated **K**inetic **E**stimation and **S**imulation) framework. Within this framework, the dynamic signaling properties of GCaMP served the dual purpose of inferring spike times and calcium handling parameters while also providing a biophysically grounded engine to bootstrap synthetic training data for high-speed ML inference.

To examine the necessity of an accurate nonlinear model, we generated ground-truth simulations of spike-evoked fluorescence using our jGCaMP8f and cell models (see **Methods**). We then performed inference using either Biophys_SMC_ or a linear autoregressive model (Linear_SMC_, adapted from ^26^) (**Supplementary Figure 7**). During elevated spiking, Linear_SMC_ produced progressively more false-positive spike predictions, particularly evident during periods of quiescent fluorescence decay (**Figure 3F, G**). Additionally, we applied Biophys_SMC_ to simulations from an explicit buffer model, which contains kinetically accurate calcium buffers along with the GCaMP model. Biophys_SMC_ accurately inferred spike times and numbers even in simulations featuring high concentrations of a parvalbumin-like calcium buffer, found commonly in GABAergic interneurons, confirming the generalizability of Biophys_SMC_ across diverse natural buffering conditions (**Supplementary Figure 8**).

Because Biophys_SMC_’s Bayesian sampling is computationally intensive, we developed Biophys_ML_ to enable fast inference at scale. We first used Biophys_SMC_ to infer cell-specific calcium handling parameters from brief (∼36 s) unlabeled fluorescence recordings using <300 s of compute time with our hardware configurations (**Methods**). We then used SMC-derived cell parameters to simulate GCaMP fluorescence driven by diverse spike trains with dataset-matched noise and baseline fluorescence drift to generate simulations matched to the kinetic properties of measured fluorescence transients. These synthetic data were used to train a supervised neural network (CASCADE ^11^ or ENS^2 12^). Once trained, Biophys_ML_ performs spike inference via a single forward pass, matching standard deep-learning runtimes while inheriting the biophysical model’s kinetics and nonlinearities (**Figure 4A**).

**Figure 4:**
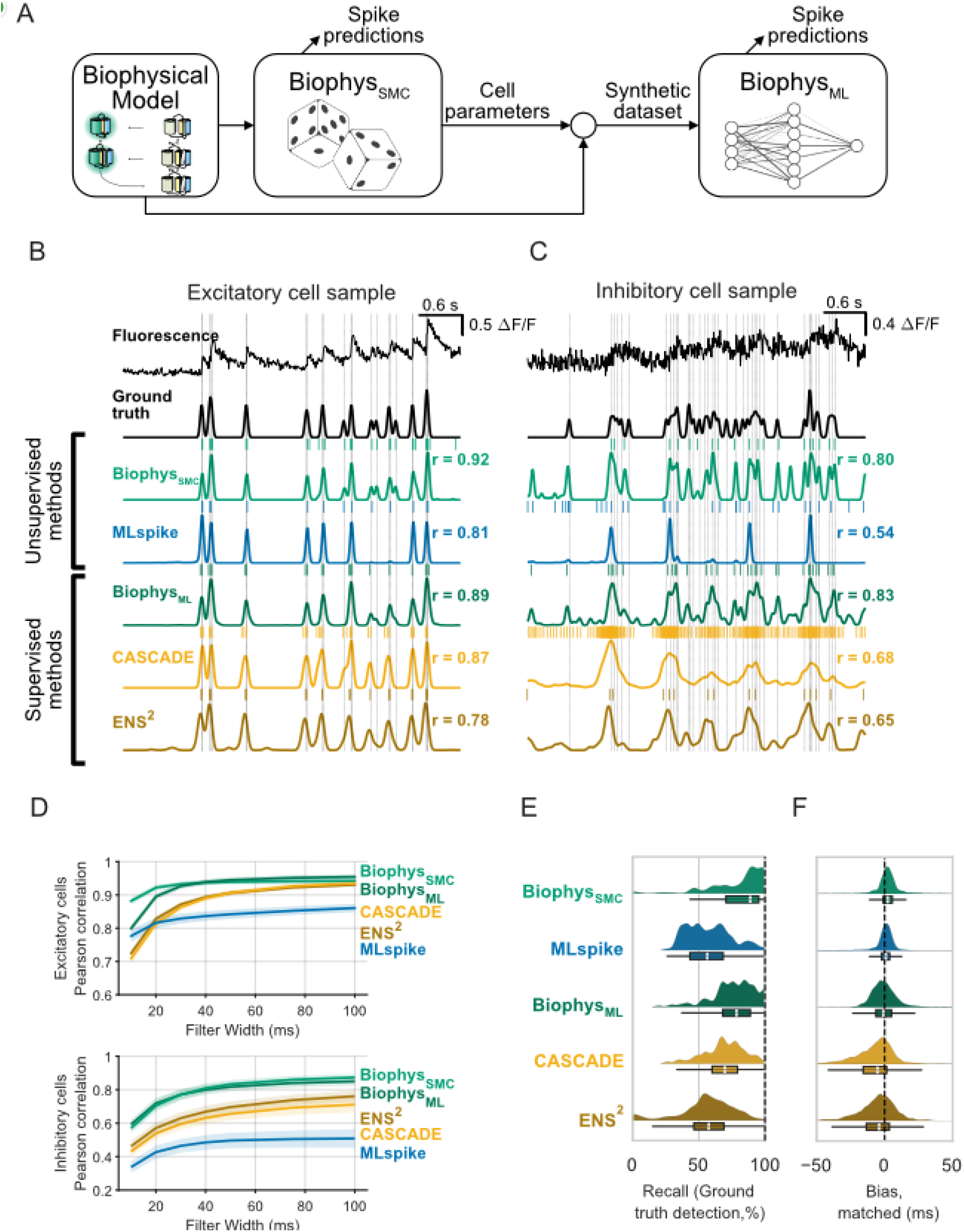
Improved action potential inference from generative model-based methods. (**A**) A generative model for imputing action potential times based on iterative fitting. The biophysical GCaMP model was held fixed while cell parameters were varied. Responses to action potential sequences were generated to fit cell parameters (BiophysSMC). After fitting was complete, the BiophysSMC model was then used to train a machine learning-based model. (**B, C**) Imputed spike probabilities using a variety of unsupervised and supervised models. Values of r indicate Pearson correlation between the time course of ground truth and that of the imputed probabilities for data filtered at 20 ms. Vertical lines indicate true spike times and tick marks indicate imputed spikes. (**B**) shows data from an excitatory cell while (**C**) shows data from an inhibitory cell. (**D**) Pearson correlations for different filtering time constants. Excitatory neurons (n = 142 epochs), mixed effects model (method and filter width as fixed effects. Method-wise comparisons: BiophysSMC vs. MLspike (p = 6.5e-62, d = 1.1), CASCADE (p = 9.0e-47, d = 0.8), and ENS2 (p = 2.3e-82, d = 1.0); BiophysML vs. MLspike (p = 3.9e-56, d = 1.1), CASCADE (p = 9.8e-42, d = 0.3), and ENS2 (p = 9.4e-76, d = 0.6). Inhibitory neurons (n = 12 epochs), mixed effects model (F(4,385) = 14.7, p = 3.6e-11). Method-wise comparisons: BiophysSMC vs. MLspike (p = 4.1e-9, d = 1.9); vs. CASCADE (p = 3.3e-4, d = 1.0); vs. ENS2 (p = 0.005, d = 1.1); BiophysML vs. MLspike (p = 1.4e-10, d = 2.0); vs. CASCADE (p = 3.3e-5, d = 1.2); vs. ENS2 (p = 8.2e-4, d = 1.4). Shaded region in each plot indicates standard error of the mean. (**E**) Discrete spike estimate statistics. Left panels show the average time between the nearest imputed and ground truth spikes, while the right shows the normalized number of spikes that were over- or underestimated. Individual points are calculated over 8-second epochs covering 1000 data points randomly selected from the full dataset. (**E-F**) Timing and detection accuracy of the different imputation models. (**E**) distribution of per-spike timing error for the various methods. Smoothed histograms and box/whisker plots summarize all errors. (**F**) epoch-wise recall of the various methods, indicating the percent of ground truth spikes recovered.

To benchmark our spike inference methods on excitatory and inhibitory cell responses from the jGCaMP8f ground truth dataset ^5^, we smoothed spike prediction traces and ground truth spikes with Gaussian filters to perform correlation analysis. Biophys_SMC_ and Biophys_ML_ predictions were tightly locked to true spike patterns for both cell classes, showing higher correlations at all Gaussian filter widths (**Figure 4B, C**). A linear mixed effects model showed that temporal correlation depended significantly on the inference method for both excitatory (F_(4,4828)_ = 135.9, p < 0.0001) and inhibitory (F_(4,408)_ = 7.2, p < 0.0001) neurons. Biophys_SMC_ and Biophys_ML_ demonstrated substantially improved continuous tracking over MLspike, CASCADE, and ENS^2^ across both excitatory (all pairwise p ≤ 0.0001, d ≥ 0.8 for Biophys_SMC_; p ≤ 0.0001, d ≥ 0.3 for BiophysML) and inhibitory datasets (all pairwise p ≤ 0.005, d ≥ 1.0 for Biophys_SMC_; p ≤ 8.2e-4, d ≥ 1.2 for Biophys_ML_) (**Figure 4D**). Training CASCADE directly on the jGCaMP8f dataset did not improve its performance (**Supplementary Figure 9**).

Next, we calculated how well the different methods were able to capture discrete spike times. To evaluate strict temporal exactness and prevent false positive predictions from artificially inflating performance metrics, we first assessed discrete spike-time capture using a narrow matching window (matching imputed spikes to the nearest ground-truth spike within a 10 ms interval; **Supplementary Figure 10**). Under these stringent criteria, our biophysical methods consistently yielded higher sensitivity (also called recall) than competing methods (**Figure 4E, Table 1**; repeated measures ANOVA, F_(4,560)_ = 91.9, p < 0.0001). Similarly, using the F-score (a metric that penalizes false positives and negatives while rewarding true positive spikes), our biophysical methods again outperformed other methods (**Table 1**; repeated measures ANOVA, F_(4,540)_ = 72.2, p <0.0001) in temporal accuracy and precision.

**Table 1.** Single-spike classification metrics for calcium-based decoding methods. Median performance across epochs for a database of 37 excitatory cells with ground-truth electrical recordings and jGCaMP8f fluorescence signals. Values reported as median (IQR). Tukey HSD post hoc results follow. Recall/Sensitivity: Biophys_SMC_ vs. MLspike p = 9.9e-9, d = 1.1; vs. CASCADE p = 9.9e-9, d = 0.6; vs. ENS^2^ p = 9.9e-9, d = 1.0; Biophys_ML_ vs. MLspike p = 9.9e-9, d = 1.1; vs. CASCADE p = 9.9e-9, d = 0.7; vs. ENS^2^ p = 9.9e-9, d = 1.0. F-score: Biophys_SMC_ vs. MLspike p = 9.9e-9, d = 1.1; vs. CASCADE p = 9.9e-9, d = 0.8; vs. ENS^2^ p = 9.9e-9, d = 0.9; Biophys_ML_ vs. MLspike p = 9.9e-9, d = 0.9; vs. CASCADE p = 9.9e-9, d = 0.6; vs. ENS^2^ p = 1.0e-8, d = 0.6. Absolute Mean bias: Biophys_SMC_ vs. MLspike p = 0.11, d = 0.2; vs. CASCADE p = 1.2e-3, d = 0.3; vs. ENS^2^ p = 1.1e-4, d = 0.4; Biophys_ML_ vs. MLspike p = 8.6e-4, d = 0.3; vs. CASCADE p = 6.3e-4, d = 0.6; vs. ENS^2^ p = 1.1e-3, d = 0.5. Uncertainty: Biophys_SMC_ vs. MLspike p = 0.01, d = 0.3; vs. CASCADE p = 1.2e-7, d = 0.5; vs. ENS^2^ p = 3.0e-7, d = 0.5; Biophys_ML_ vs. MLspike p = 0.6, d = 0.1; vs. CASCADE p = 1.2e-7, d = 0.5; vs. ENS^2^ p = 2.9e-5, d = 0.4.

|  | Precision | Recall | F-Score | Accuracy | Mean bias (ms) | Uncertainty (ms) |
| --- | --- | --- | --- | --- | --- | --- |
| Biophys<sub>SMC</sub> | 0.84 (0.75, 0.92) | 0.88 (0.71, 0.95) | 0.83 (0.71, 0.91) | 0.71 (0.55, 0.83) | 1.9 (-2.9, 3.0) | 3.8 (1.8, 7.3) |
| MLspike | 0.70 (0.53, 0.93) | 0.56 (0.43, 0.68) | 0.60 (0.51, 0.70) | 0.43 (0.34, 0.54) | -2.0 (-0.6, 0.9) | 3.2 (1.5, 6.2) |
| Biophys<sub>ML</sub> | 0.76 (0.68, 0.84) | 0.78 (0.68, 0.89) | 0.75 (0.70, 0.80) | 0.61 (0.54, 0.67) | 0.0 (-2.9, 3.0) | 5.8 (2.7, 10.3) |
| CASCADE | 0.69 (0.63, 0.78) | 0.70 (0.60, 0.79) | 0.69 (0.61, 0.74) | 0.52 (0.45, 0.59) | -7.4 (-11.2, -3.2) | 8.2 (3.5, 18.2) |
| ENS<sup>2</sup> | 0.71 (0.65, 0.86) | 0.57 (0.46, 0.69) | 0.65 (0.56, 0.74) | 0.48 (0.39, 0.58) | -6.6 (-12.3, -2.3) | 8.3 (3.8, 16.5) |

While a narrow band rigorously tests exactness, it penalizes near-miss predictions without capturing their underlying error distribution. To determine whether false positives were random artifacts or systematic shifts, we evaluated bias and uncertainty within a wider 100 ms interval (**Figure 4F, Supplementary Figure 11**). This expanded window revealed a systematic bias in which predicted spikes preceded ground-truth times for both supervised methods (CASCADE, 7.45 ms; ENS2, 6.61 ms) and MLspike (2.02 ms). In contrast, the biophysical methods successfully reduced this bias (**Table 1**; repeated measures ANOVA, F_(4,556)_ = 13.0, p <0.0001) and reported spike times with significantly lower uncertainty and superior epoch-wise statistics (**Table 1**; repeated measures ANOVA, F_(4,556)_ = 16.2, p<0.0001).

To assess the generalizability of our biophysical approach, we evaluated its robustness across varying sampling rates, training datasets, and calcium indicators. First, we applied our inference methods alongside CASCADE, ENS2, and MLspike to downsampled versions of the Janelia jGCaMP8f dataset (10 and 30 Hz, rates commonly used in two-photon imaging). Our biophysical methods consistently outperformed the others in correlation to ground truth across all tested frequencies (**Supplementary Figure 12**; Linear mixed effects model, fixed effects of method, F_(4,1988)_ = 83.6, p<0.0001).

Next, we examined whether external *in vivo* data could substitute for our *ex vivo* slice recordings to estimate GCaMP model parameters. By constraining the jGCaMP8f model using subsets of the Janelia dataset containing both high and low activity periods (see **Methods**), we achieved statistically equivalent spike inference performance with Biophys_SMC_ (**Supplementary Figure 13**). This indicates that our model can be effectively constrained using any ground truth dataset featuring sufficient dynamic range to engage nonlinear sensor properties.

Finally, we tested the framework’s adaptability to other indicators. We found that the slower indicator, jGCaMP8m, also exhibited activity-dependent decay slowing (**Supplementary Figure 14A-C**, Linear mixed effects model, fixed effect of *τ*_1_; F_(1,162.4)_ = 177.8, *β* = 0.085, p<0.0001). After fitting the GCaMP model using stopped-flow and *in vivo* jGCaMP8m Janelia recordings (**Supplementary Figure 14D, E**), our biophysical methods again outperformed CASCADE, ENS2, and MLspike (**Supplementary Figure 14F, G**, Linear mixed effects model, main effect from method; F_(4,6596)_ = 185.8, p<0.0001). Taken together, these results demonstrate that our C-SPIKES inference framework generalizes robustly across diverse imaging conditions, datasets, and sensor variants.

**Supplementary Figure 9:**
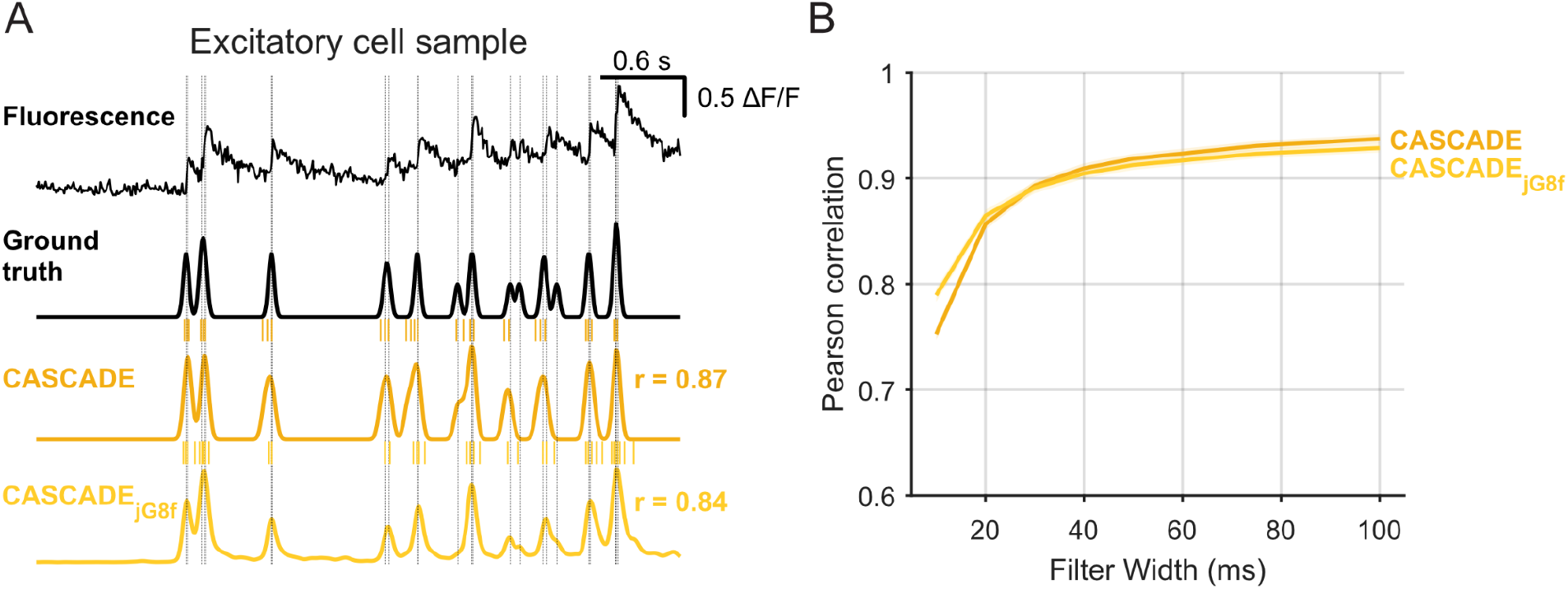
Training on jGCaMP8f data does not improve CASCADE performance. (**A**) Imputed spike probabilities using the CASCADE Global Exc model or one trained on the jGCaMP8f dataset itself. Values of r indicate Pearson correlation between the time course of ground truth and that of the imputed probabilities for data filtered at 20 ms. Vertical lines indicate true spike times and tick marks indicate imputed spikes. (**B**) Pearson correlations for different filtering time constants.

**Supplementary Figure 10:**
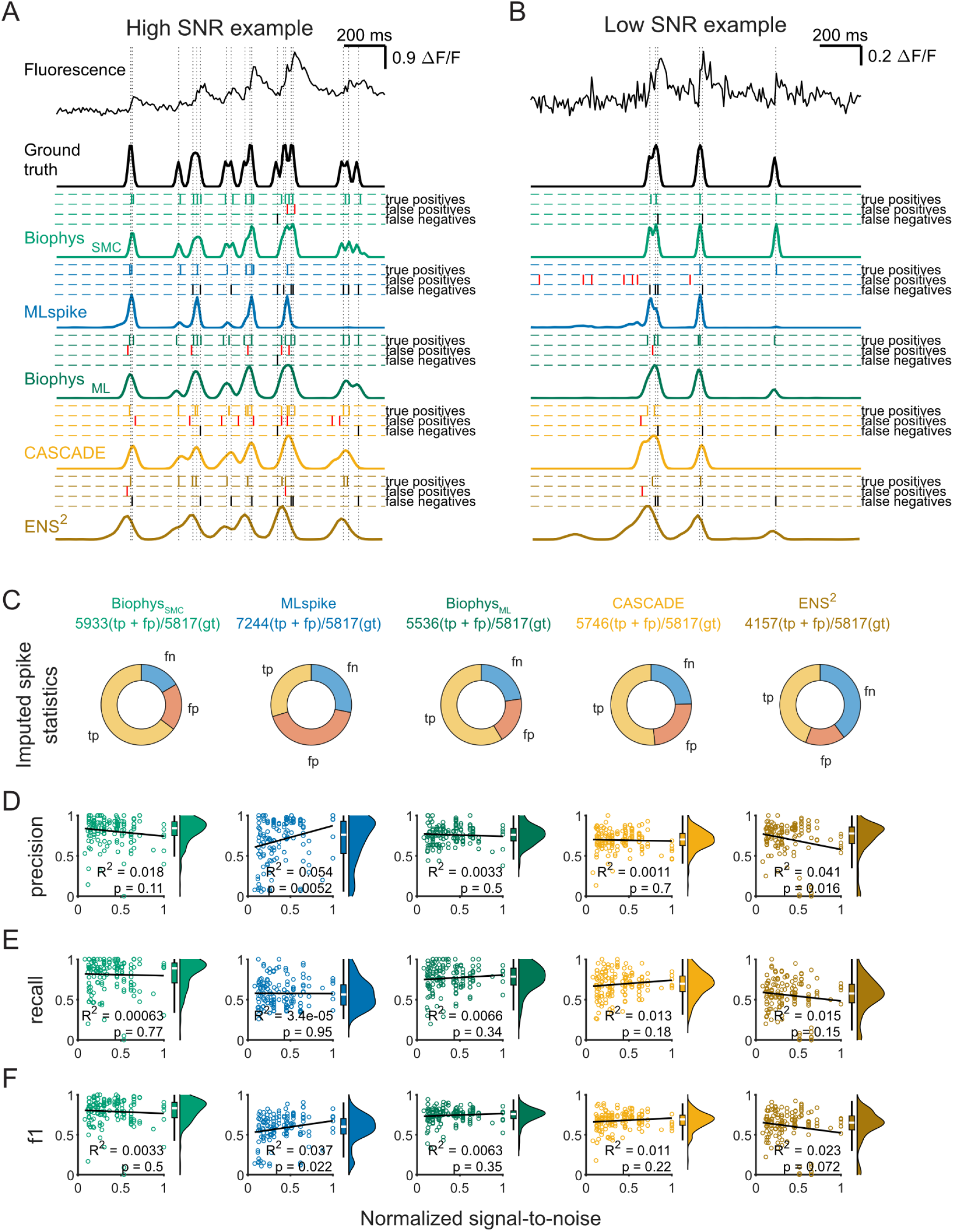
Statistics of discrete spike estimate comparisons to ground truth spike times. (**A, B**) Continuous and discrete spike predictions across methods. The top two traces (black) show raw fluorescence and Gaussian-convolved (10 ms) ground truth spike times. Ground truth spike times are indicated as vertical tick marks. Discrete imputed spikes for each method are shown above the corresponding continuous prediction trace, sorted into true positives, false positives, and false negatives. An epoch was selected from a representative cell which produced high signal-to-noise ratio (SNR) fluorescence changes in response to an action potential (**A**) or low SNR (**B**). (**C**) Columns show the proportions of imputed spikes that were scored as true positive (tp), false positive (fp), or false negative (fn) for each method with a comparison of all imputed spikes produced by each method (tp + fp) compared to total number of ground truth (gt) spikes from the 37 putative excitatory cells in the Janelia dataset. (**D-F**) Precision (tp/(tp + fp), **D**), Recall (tp/(tp + fn), **E**), and F-score (harmonic mean of recall and precision, **F**) as a function of the SNR of each recorded epoch. The black line in each plot indicates a regression fit to the data with r^2^ indicating the amount of variation explained by SNR and p indicating significance of the t-statistic comparing a linear to a constant model. Of all methods, only MLspike shows reduced performance at the SNR levels tested across this dataset. In particular, the precision (propensity to find false negatives) was increased in low SNR conditions.

**Supplementary Figure 11:**
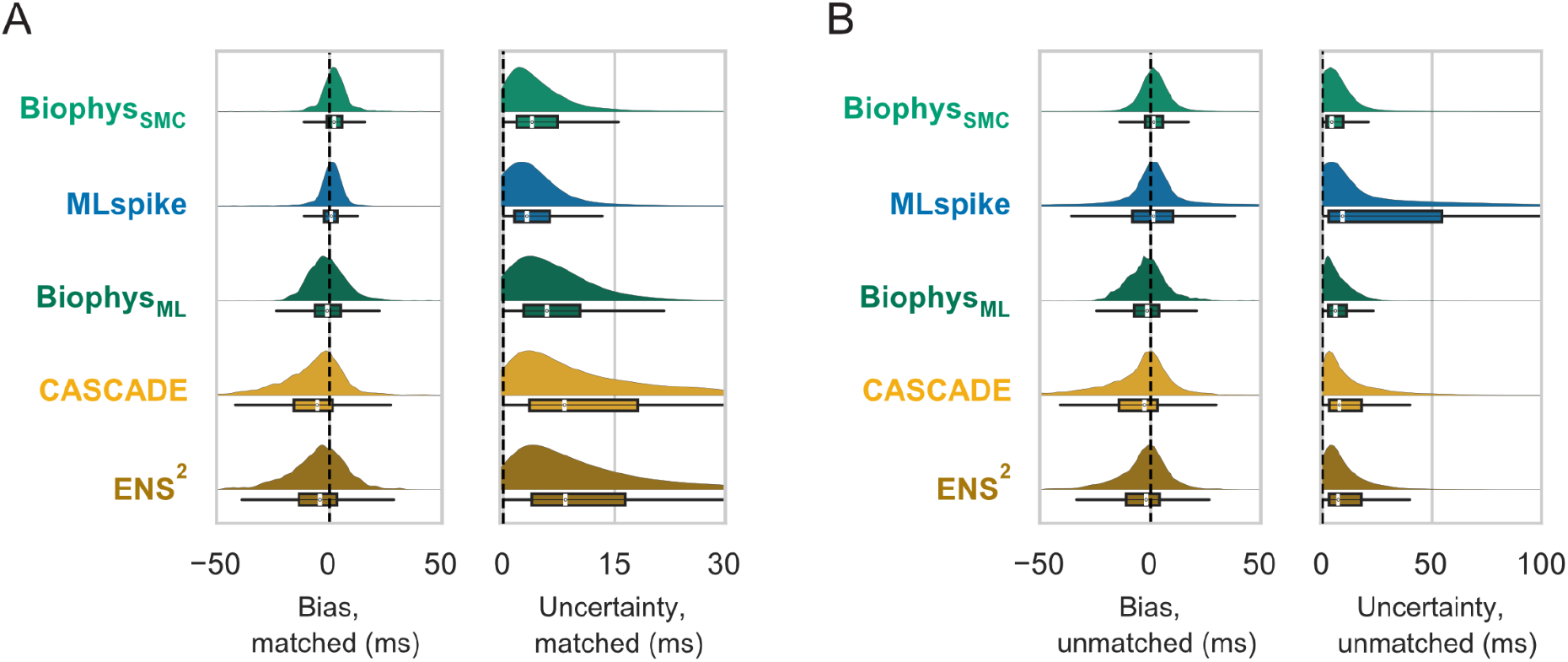
Comparison of matched and unmatched temporal differences across methods. (**A**) Bias and uncertainty between imputed and ground truth spikes matched within a 100 ms window. Each plot is a kernel density estimate over a boxplot of indicated metric for each imputed spike. (**B**) As **A** for all imputed spikes compared to the nearest ground truth spike. Note that the supervised methods show increased bias and uncertainty when removing double-counted imputed spikes by using matched statistics.

**Supplementary Figure 12:**
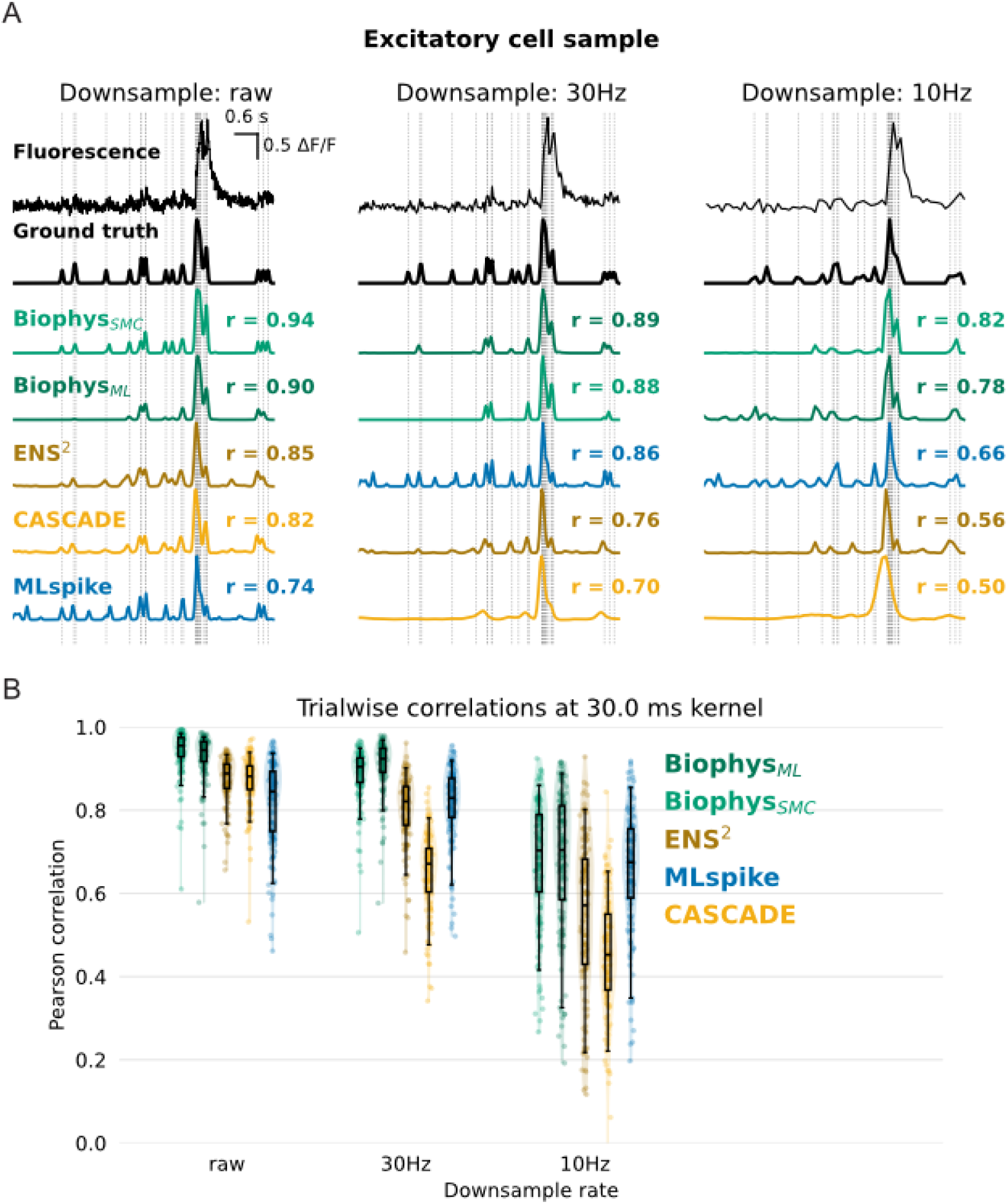
Impact of acquisition rate on spike inference performance. (**A**) Sample fluorescence trace (top) with raw (left), 30 Hz (middle), and 10 Hz (right) mean downsampling applied to demonstrate how performance is impacted by simulated acquisition rate. In each panel, ground truth traces convolved to a 30 ms Gaussian filter on the corresponding downsampled time base (second to top) are followed by predictions using the various inference methods convolved by the same filter with Pearson correlation shown for each prediction at trace left. (**B**) Distributions of correlations with ground truth achieved with each method as a function of acquisition downsampling rate at 30 ms Gaussian filter. Each point corresponds to a single epoch. Mixed-effects modeling indicated a strong effect of method on correlation across smoothing conditions. In smoothing-specific comparisons, both biophysical methods consistently outperformed CASCADE and ENS^2 at all smoothing levels (all p <= 1.1e-17; d = 0.88-3.64). Relative to MLspike, both biophysical methods outperformed it at 30 Hz and raw smoothing (all p <= 1.3e-46; d = 1.00-1.22), whereas at 10 Hz the difference was small and not significant (BiophysSMC: p = 0.75, d = 0.12; BiophysML: p = 0.13, d = 0.12) with this null result likely accounted for by false positives produced by MLspike reported throughout this work.

**Supplementary Figure 13:**
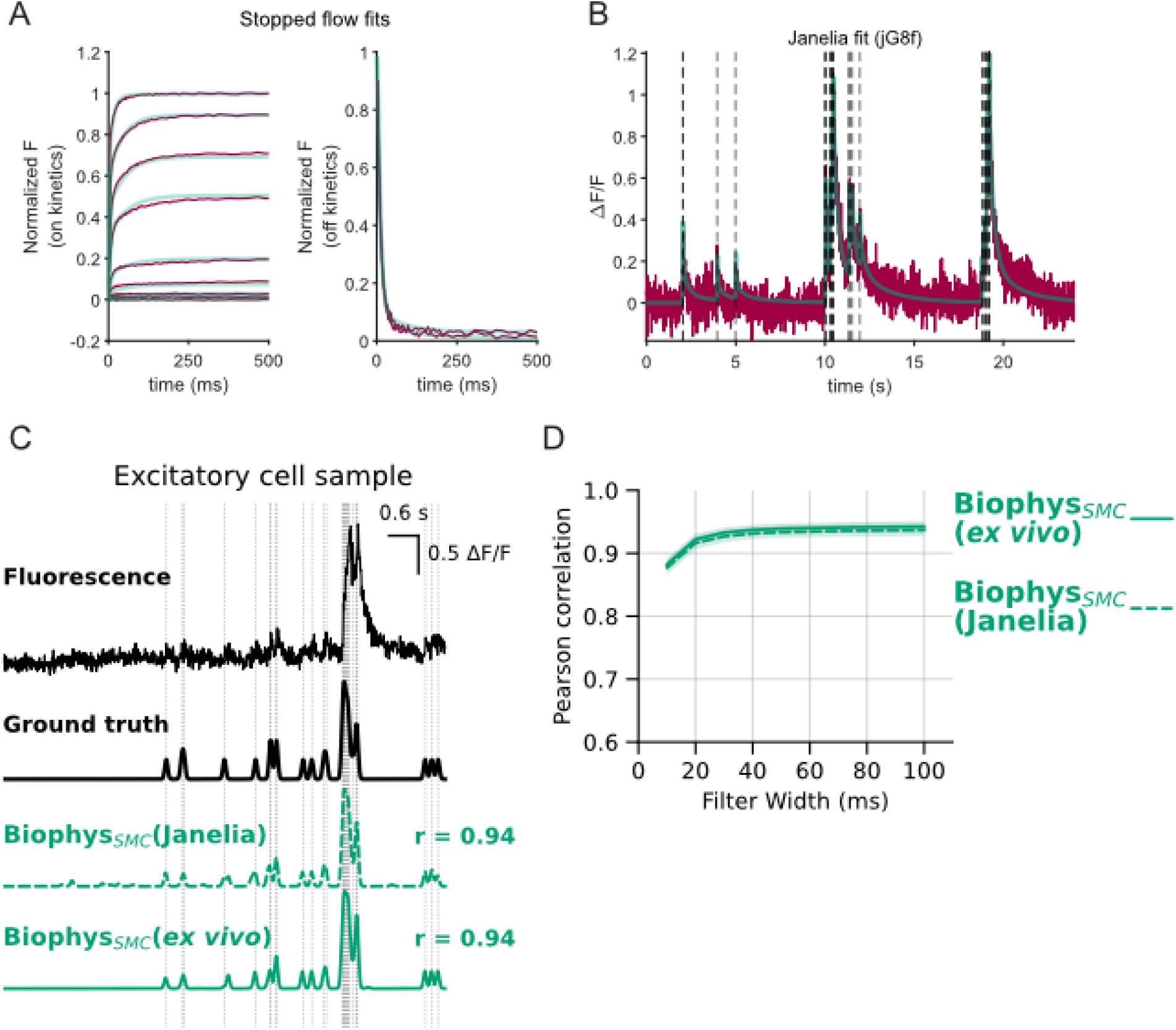
Generalization of ground truth source for jGCaMP8f biophysical model parameter selection. **(A)** Global GCaMP model fit to the jGCaMP8f stopped flow data for the selected GCaMP model (see **Methods**). (**B**) Fit of selected GCaMP model run through the generative cell model to a sample of the Janelia jGCaMP8f dataset. Unlike in Supplementary Figure 5, here, the GCaMP model was selected on the basis of its ability to fit the Janelia dataset itself. (**C**) Sample fluorescence trace (top) with ground truth spikes as dashed vertical lines and expressed as a trace by convolution with a 30 ms Gaussian kernel (second to top). Predictions made by Biophys_SMC_ using GCaMP parameters constrained by the Janelia dataset (second to bottom) or our ex vivo dataset (bottom) are shown convolved by the same kernel width. The Pearson’s correlation of the prediction and ground truth traces is shown right of each prediction trace. (**D**) Median correlations between the ground truth and prediction traces convolved with kernels of increasing filter width. The correlations are statistically indistinguishable between the Biophys_SMC_ predictions made using GCaMP parameters selected using either ground truth data source.

**Supplementary Figure 14:**
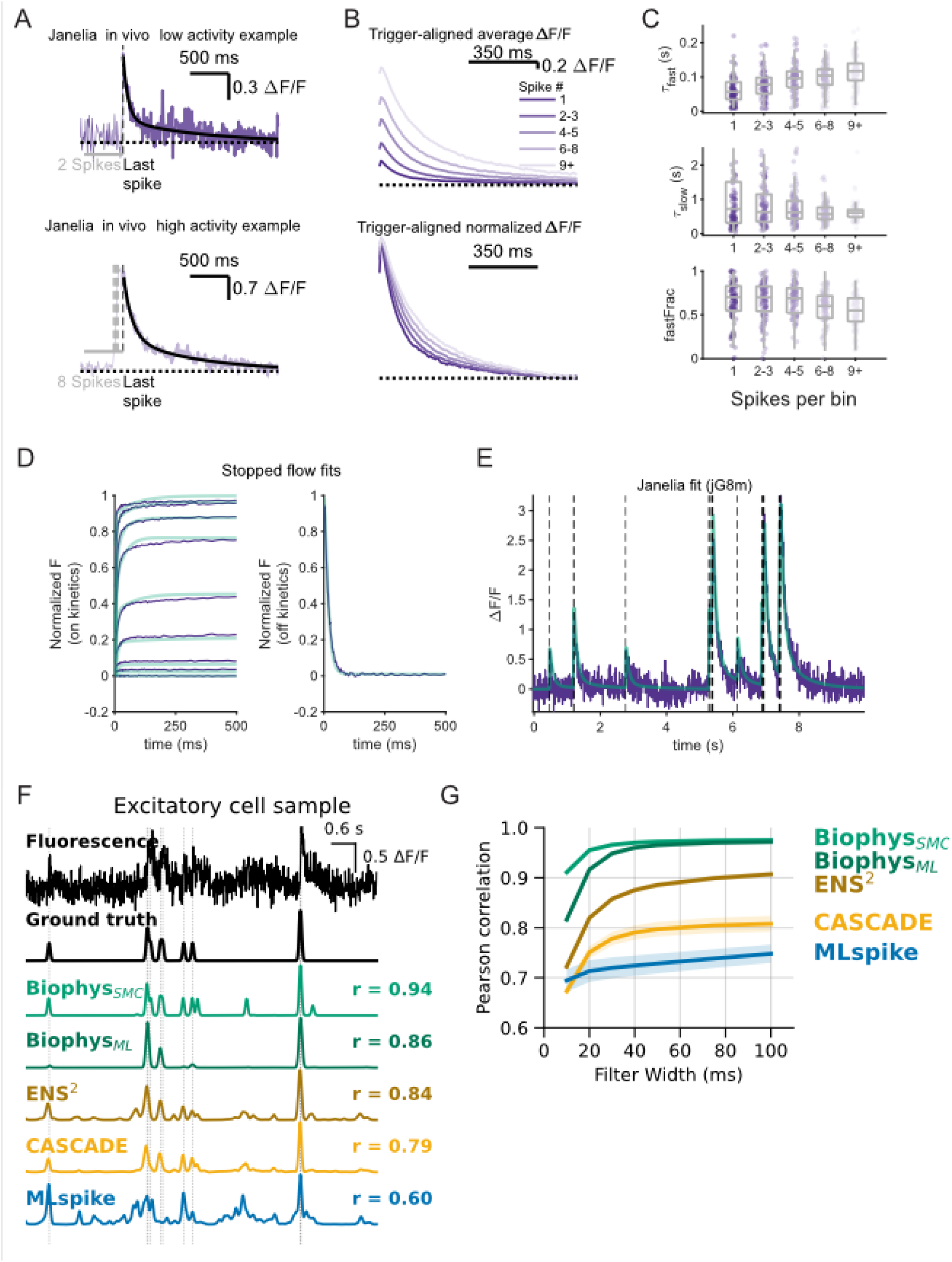
Development and performance benchmarking of biophysical methods for jGCaMP8m. (**A-C**) Activity-dependent elongation of fluorescence decay generalized to jGCaMP8m responses in vivo. (**A**) Examples of the fluorescence tail following low or high numbers of spikes. **(B)** Aligning of fluorescence response by binned spike number in the 500 ms preceding the period shown. The bottom panel shows the alignment of normalized progressively higher activity demonstrating the in vivo elongation effect. (**C**) Boxplot of fit fast time constant (top), slow time constant (middle), and contribution of each (bottom) to the decay of individual events binned by spike number in the count window. Mixed effects model included 2654 observations from n = 194 epochs. (**D**) Global GCaMP model fit to the jGCaMP8m stopped flow data for the selected GCaMP model (see Methods). (**E**) Fit of the cell-specific parameters (using the fixed GCaMP model) to a sample recording from the Janelia jGCaMP8m dataset. (**F**) Sample fluorescence trace (top) with ground truth spikes as dashed vertical lines and expressed as a trace by convolution with a 30 ms Gaussian kernel (second to top). Predictions made by Biophys_SMC_ using GCaMP parameters constrained by the Janelia dataset (second to bottom) or our ex vivo dataset (bottom) are shown convolved by the same kernel width. The Pearson correlation of the prediction and ground truth traces is shown right of each prediction trace. (**G**) Median correlations between the ground truth and prediction traces convolved with kernels of increasing filter width. Mixed effects model shows that correlation depends on the method used (method and kernel width as fixed effects, F(4, 6596) = 185.8, p = 4.1e-151). Method-wise comparisons demonstrate that our methods outperform MLspike (BiophysSMC: p < 2.2e-308, d = 1.6; BiophysML: p < 2.2e-308, d = 1.3), CASCADE (BiophysSMC: p < 2.2e-308, d = 1.6; BiophysML: p < 2.2e-308, d = 1.3), and ENS2 (BiophysSMC: p < 2.2e-308, d = 1.8; BiophysML: p < 2.2e-308, d = 1.5).

## Discussion

The complex, nonlinear kinetics of fluorescent indicators impact accurate spike inference in ways that have seldom been explored. Here, we developed C-SPIKES, a biophysically-inspired generative approach to explicitly account for the sensor-imposed distortions to the spike-to-fluorescence transformation. By characterizing the temporal distortions imposed by GCaMP, we parameterized a generative model which became the basis for an SMC-based approach to spike detection (Biophys_SMC_) which also serves as the basis for synthetic training data generation. This strategy allowed us to train a supervised network (Biophys_ML_) that achieves the computational speed of current deep-learning methods while delivering the superior accuracy and generalizability of detailed biophysical models, fundamentally decoupling high-performance spike inference from the need for laborious experimental ground-truth datasets.

In the present work, we first characterized the underlying nonlinear distortions in the spike-to-fluorescence signals of four fast GCaMP variants (GCaMP6f, jGCaMP7f, jGCaMP8f, and jGCaMP8m). While earlier variants (6f and 7f) exhibited a facilitation-like nonlinearity in their fluorescence response to closely-timed spikes (also noted in ^10,27–29^), which last tens of milliseconds. We also report the discovery of a novel, persistent use-dependence in all tested GCaMPs that progressively slows fluorescence decay by up to three-fold during sustained activity. Inspired by the known kinetics of calcium binding to calmodulin ^33^, we propose that this use-dependent slowing arises from the kinetic asymmetry between the fast N-lobe and slow C-lobe. Specifically, the C-lobe acts as a high-affinity reservoir that sequesters calcium, effectively trapping the sensor in a slow-decaying, non-fluorescent state during high-frequency stimulation.

GCaMP’s use-dependent slowing violates the linear spike-to-fluorescence assumptions of many existing decoders. Consistent with this framing, a linear SMC approach generated spurious spike predictions as the elongated decay dynamics forced latent states to improperly match the fluorescence (**Figure 3G, H**). Further, MLspike, which uses a linear generative model, performed poorly under high-spike-rate, low-SNR conditions (**Figure 4C; Supplementary Figure 9C**).

To parametrize our proposed GCaMP model kinetics independently from cell-intrinsic calcium handling, we combined *in vitro* calcium clamp measurements with *ex vivo* spike-dependent fluorescence measurements. We first used stopped-flow and static fluorimetry on purified protein, where the calcium step input and ionic and buffer solutions are precisely defined. The resulting calcium-dependent kinetics and fluorescence amplitudes were used to constrain model calcium-dependent transition rates while steady-state fluorimetry measurements constrained the maximum-to-minimum fluorescence ratio. We then used acute-slice recordings in a preparation with well-characterized calcium buffering and linear presynaptic calcium entry per spike ^30^ to unambiguously identify indicator nonlinearities (e.g., supralinear summation and activity-dependent slowing during spike trains).

We used a fixed GCaMP model embedded within our cell model to create Biophys_SMC_, an unsupervised sequential Monte Carlo sampler. Biophys_SMC_ infers spikes with high reliability while simultaneously estimating cell-specific parameters (including calcium-handling and spiking statistics). This approach yielded the highest accuracy in spike inference. Furthermore, regardless of the chosen binning resolution, this generative approach provides a distinct advantage over supervised methodologies by naturally expressing the posterior distribution of spike counts as a probability with an associated uncertainty.

As we show, Monte Carlo methods can yield highly accurate posterior state predictions ^26^, however, they incur a high computational cost ^13,38^. Conversely, deep learning approaches like ENS^2^ and CASCADE offer rapid inference, but they rely on large labor-intensive recordings of ground-truth datasets and as we show, they can fail to generalize to sensors with properties not present in the training dataset. Indeed, both ENS^2^ and CASCADE exhibited temporal biases and overestimated rise times for jGCaMP8f, highlighting the fragility of networks trained on broad datasets (**Figure 4, Supplementary Figure 13**).

We addressed the tradeoff between accuracy, computational cost, and data availability in two ways. First, we implemented a GPU-acceleration for Biophys_SMC_ to partially mitigate the computational cost of the method (an approximate 600x speed up compared to a single thread CPU implementation). Then, we used this accelerated Biophys_SMC_ to create our novel generative machine-learning pipeline. By forward-simulating fluorescence traces based on the inferred cell-specific parameters, our model generates virtually unlimited, fit-for-purpose synthetic training data.

This synthetic data was used to train Biophys_ML_, which performs inference as rapidly as CASCADE or ENS2 but with substantially higher accuracy and robustness. Crucially, this generative approach allows a limited cell- or experiment-specific (e.g., anesthetized vs. awake) dataset to provide estimates of the posterior distributions needed to generate a training dataset spanning a wide range of spike timing patterns. To our knowledge, our study is the first to report improvements in machine learning training by using a generative modeling approach.

In summary, biophysically constrained generative modeling offers a powerful, extensible strategy for estimating spike probability from nonlinear sensor signals. By separating sensor characterization from downstream inference, the C-SPIKES framework offers out-of-the-box performance and overcomes the data-dependency bottlenecks of current deep learning approaches. The strategies introduced here should readily generalize to emerging biosensors, including red-shifted indicators ^39–41^, chemigenetic sensors ^42,43^, and reporters of membrane voltage ^44,45^ or neuromodulator release ^46,47^, providing a unified path toward mechanistically informed decoding across diverse optical tools.

## Methods

### Animals

The following strains were used for physiological experiments: C57Bl/6J (Jackson Laboratories stock 000664); CB6F1 (BalbC and C57Bl/6J F1); and Gabra6 (B6;129P2-Gabra6tm2(cre)Wwis/Mmucd, MMRRC stock 000213-UCD). Both males and females were used and were between 40 and 130 days old. Animals were used in accordance with Princeton University’s Institutional Animal Care and Use Committee and Institut Pasteur’s CEEA-Paris1 approved protocols.

### Protein purification

GCaMP variants were cloned into the pET28b protein expression vector using GCaMP6f, 7f, and 7s backbones generously provided by Douglas Kim (Janelia) and Addgene (104488). Plasmids were transformed into IPTG-inducible BL21(DE3) competent E. coli (MilliporeSigma 694504). Starter cultures of transformed cells were grown in 5 ml LB medium with 50 mg/L kanamycin at 37°C for 4 hours. Cultures were then added to 1 L LB medium with 50 mg/L kanamycin and grown at 37°C until the OD at 600 nm reached 1.0. 1 mM IPTG was added and the incubation temperature dropped to 25°C overnight for optimal protein expression. Liquid cultures were centrifuged at 6,000 relative centrifugal force (rcf) for 10 minutes.

Pellets were resuspended in 25 mM Tris-HCl, 500 mM NaCl, 20 mM imidazole, pH 8, with 1 mM PMSF for protease inhibition. Cells were mechanically lysed through an Emulsiflex homogenizer (ATA Scientific). Cell debris was removed through centrifugation at 13,000 rcf for 45 minutes at 4°C and the supernatant bound to nickel-NTA agarose (ThermoScientific 25215). Protein was eluted with 25 mM Tris-HCl, 500 mM NaCl, 500 mM imidazole, pH 8. Proteins were further concentrated using 10 kDa MWCO centrifugal filters (MilliporeSigma UFC901008) and desalted (GE Healthcare 17085101) into 130 mM KMOPS (in mM:100 KCl, 30 MOPS, pH 7.2), or a buffer mimicking intracellular conditions (in mM: 125 KCl, 14.5 KOH, 30 NaCl, 0.7 MgCl2, 10 HEPES) termed internal buffer as modified from ^48^. Protein concentrations were determined by absorption spectrophotometry using alkali denaturation in 0.1M NaOH. Under these conditions, mature GCaMP chromophore exhibits a standard molar absorption coefficient of 44,000M-1cm-1 at 447 nm, allowing direct recovery of GCaMP concentration using Beer’s law ^49,50^.

### Protein steady-state and kinetic characterization

Excitation and emission spectra were measured on a PTI Quantamaster 800 spectro-fluorometer (Horiba Jobin Yvon Inc., Edison, NJ, USA) with data collection under control of Felix GX software. For calcium-dependent fluorescence, steady-state measurements were made at 20°C using 0.15 μM of purified protein suspended in zero-Ca2+ buffer or high-Ca2+ buffer from a calcium calibration kit (ThermoFisher C3008MP). The zero-Ca2+ buffer was reciprocally diluted with the high-Ca2+ buffer to reach free Ca2+ concentrations between 0.01 and 10 μM. Hill coefficient and Kd were estimated by fitting steady state data to the Hill equation.

The fluorometer was also used to prepare reagents for stopped-flow kinetic measurements. Nominally Ca2+-free buffers included KMOPS or internal buffer supplemented by 2 mM BAPTA (Invitrogen B1204). Increasing amounts of 1 M CaCl2 were added to create free calcium steps of 0 nM, 50 nM, and increasing by approximately 2x free calcium in ten steps to 10 μM free calcium. Indicators were diluted to 175 nM in 0.1mM solution buffered by 0.1 mM BAPTA (see design considerations in **Supplementary Figure 2**).

Off-response kinetics buffers were designed to have a starting free calcium concentration of 10 μM in 100 μM BAPTA and a final nominal concentration of 0 nM or 50 nM free calcium when mixed to Ca2+-free buffer. Proteins were diluted to 35 nM in the 10 μM free calcium buffer. In all cases, fura-2 (Molecular Probes F6799) and fura-4F (Invitrogen F14174) were used to calibrate actual free calcium concentrations.

On-response and off-response kinetics were measured using an Applied Photophysics SX20 stopped-flow spectrometer. Excitation light was provided by a xenon arc lamp directed through a monochromator set to a 4 nm passband centered on 480 nm, while emissions were collected through a 550 nm longpass filter (Newport FR-OG550). Dead time in the range of 0.6-1.5 ms was estimated by extrapolation from measurements using OGB-1 (Life Technologies, O6806). Measurements were made at 37°C with 60 µl per reaction chamber.

The success of biophysical modeling depended on characterizing sensor kinetics under conditions that rigorously mimicked the intracellular environment. We found that jGCaMP8f (and the other jGCaMP8 variants, unpublished observations) is significantly more sensitive to competitive binding by magnesium than previous generations (Supplementary Figure 3). Standard characterizations performed in magnesium-free buffers would therefore yield kinetic constants and photophysical properties that fail to predict in vivo performance. Furthermore, we demonstrated that the design of the calcium clamp itself (specifically the buffering capacity relative to sensor concentration) can distort measured off-rates if not carefully controlled (**Supplementary Figure 2**). Had we relied on standard stopped-flow protocols without these corrections, the resulting model would have failed to reconcile the kinetic and imaging datasets, rendering the subsequent generative modeling invalid.

### Adeno-associated virus constructs

Adeno-associated viruses (AAVs) were prepared by cloning the GCaMP variants into pAAV backbones (GenScript, Piscataway, NJ). Plasmids were incorporated into AAV serotype 1 (AAV2/1) and serotype DJ (AAV DJ) by the Princeton Neuroscience Institute Viral Neuroengineering Laboratory.

### Stereotactic viral Injection

Viral vectors were injected stereotactically into the cerebellar vermis. Mice were deeply anesthetized by intraperitoneal injections of 1.5% ketamine (Mérial) and 0.05% xylazine (Bayer). 2% Xylocaine (Newpharma) was applied to the cranial incision. 100 nl vermal (6.5 mm caudal to bregma, lateral 0.2 mm, ventral 3.6 mm and 3.4 mm) and 100 nl Crus I (6 mm caudal to bregma, lateral 3 mm, ventral 0.5 mm) injections were infused over one minute. To allow for optimal transgene expression, mice were imaged 2 to 4 weeks after injection.

### Slice preparation

Animals were euthanized by rapid decapitation. The brains were quickly removed and placed in an ice-cold solution containing (in mM) 2.5 KCl, 0.5 CaCl2, 4 MgCl2, 1.25 NaH2PO4, 24 NaHCO3, 25 glucose, 230 sucrose, and 0.5 ascorbic acid. The buffer was bubbled with 95% O2 and 5% CO2. A Leica VT1200S vibratome was used to prepare coronal slices (200 µm), which were incubated at 32 °C for 30 min in (in mM): 85 NaCl, 2.5 KCl, 0.5 CaCl2, 4 MgCl2, 1.25 NaH2PO4, 24 NaHCO3, 25 glucose, 75 sucrose, and 0.5 ascorbic acid. The external recording solution contained (in mM) 125 NaCl, 2.5 KCl, 1.5 or 2.0 CaCl2, 1.5 or 1.0 MgCl2, 1.25 NaH2PO4, 25 NaHCO3, 25 glucose, and 0.5 ascorbic acid. Slices were maintained at room temperature for up to 6 h. All slice recordings were performed at 36–38°C.

### Imaging in cerebellar brain slices

Parallel fiber boutons expressing fluorescent virus were imaged with an Ultima two-photon scanning scanhead (Bruker Nano Surfaces Division, Middleton, WI, USA) that was mounted on an Olympus BX61W1 microscope equipped with a water-immersion objective (60×/1.1-NA; Olympus Optical, Tokyo, Japan) and infrared Dodt-gradient contrast. Two-photon excitation was performed with a pulsed Ti:Sapphire laser (DeepSee, Spectra-Physics, France) tuned to 920 nm. Extracellular parallel fiber axonal stimulation was performed with a constant voltage stimulator (Digitimer Ltd, Letchworth Garden City, UK) and a patch pipette (typically with a tip resistance of 4–6 MΩ) filled with artificial cerebrospinal fluid and placed in the molecular layer adjacent to labeled parallel fibers. The fluorescence of single cerebellar granule cells was monitored using circular line scans along the cytoplasm of the GC somata or straight line scans across individual parallel fiber boutons. Typical line rates were ∼1 kHz. To determine threshold fluorescence, extracellular stimulation intensity was increased until clear fluorescence responses were observed after a train of 20–60 60-μs pulses at 100 or 300 Hz. To ensure action potential initiation for each stimulus pulse, the stimulus voltage was increased by 5 V above threshold.

### Biophysical and cell models

#### Linear kernel fitting

For modeling linear bouton responses in **Figure 1N, O**, fluorescence traces were modeled as a linear time-invariant system driven by the spike train. We denote the normalized fluorescence as *y*(*t*) = Δ*F* / *F* (*t*) and the spike train as 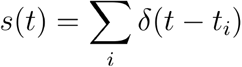 with the predicted response given a convolution with impulse response *h*, 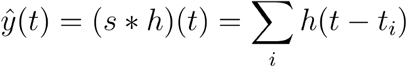. This impulse response was parameterized as a three-parameter rise-decay kernel,

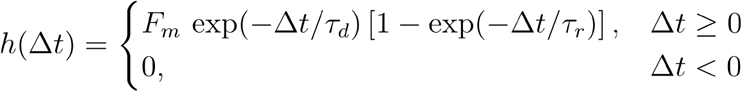

where *τ*_*r*_ and *τ*_*d*_ are rise and decay time constants and *F*_*m*_ is an amplitude scale factor.

Parameters (*τ*_*d*_ *τ*_*r*_ *F*_*m*_) were estimated by constrained nonlinear least-squares minimization (*lsqcurvefit*, MATLAB) using the measured single-spike response after stimulus-time alignment and baseline-window truncation. The fitted kernel was then used without further adjustment to predict responses to multi-spike protocols (paired pulses and Poisson spike trains) by linear superposition.

#### GCaMP model

We modeled GCaMP fluorescence as arising from semi-independent (“hemi-independent”) ^36^ calcium binding at two lobes, each of which can subsequently bind a target peptide to become fluorescent. Formally, each lobe transitions between an unbound and a calcium-bound state, which is governed by first-order binding and unbinding reactions:

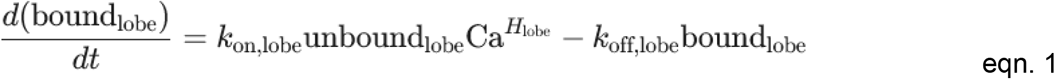

where *k*_on,lobe_ and *k*_off,lobe_ are rate constants for binding and unbinding at each lobe while Ca represents the free calcium and *H*_lobe_ is a Hill coefficient describing cooperative binding. We defined:

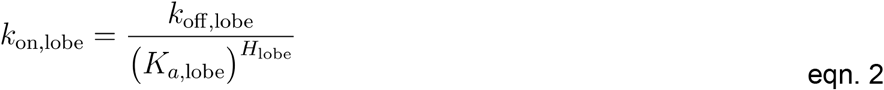

where *K*_*a*,lobe_ is an apparent dissociation constant. Once a lobe is bound with calcium, it can transition to a fluorescent state by binding the target peptide. This step similarly follows first-order kinetics:

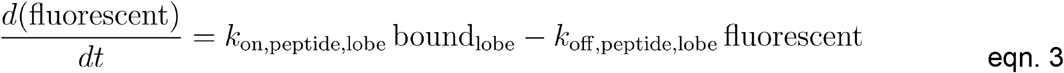

All fluorescent states (i.e., those in which a lobe is calcium-bound and has the peptide bound) contribute equally to total fluorescence. For stopped-flow simulations, model fluorescence was normalized between 0 (occupancy of fluorescent states at zero calcium) and 1 (occupancy at saturating calcium). For cell-based simulations, fluorescence was scaled to a dimensionless quantity *F* by setting it to 1 at 0 calcium and the parameter *R*_*f*_ – the dynamic range of the indicator – at saturating calcium level. 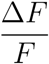 was then calculated as:

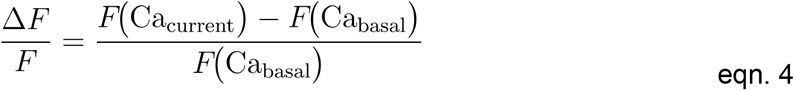

where *F* (Ca_current_) is the instantaneous normalized fluorescence at the current free-calcium level, and *F* (Ca_basal_) is the steady-state fluorescence at basal calcium.

#### Explicit Buffer model

For simulated traces presented in **Supplementary Figure 8**, the “jGCaMP8f Slice” GCaMP model (**Table 4**) was exposed to a single compartment calcium model (Rebola et al., 2019) where intracellular calcium levels follow a differential equation of the form:

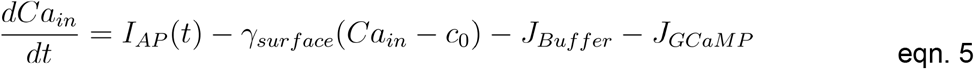

where *I*_AP_ (*t*) is the calcium influx due to a spike modeled as the sum of Gaussian kernels centered at a specified offset from each spike time with integral Δ*Ca*_*spike*_. The term *γ*_*surface*_ (*Ca*_*in*_ − *c*_0_) accounts for extrusion across the plasma membrane, and *J*_*Buffer*_ and *J*_*GCaMP*_ represent the action of calcium binding and unbinding buffers (endogenous buffer, ATP, and Parvalbumin) or GCaMP, respectively.

The binding kinetics of GCaMP were modeled as *J*_*GCaMP*_ to account for the buffering action of the sensor itself. To accurately reflect the cooperative binding architecture of the indicator described above, this term represents the net rate of calcium binding summed across the individual modeled domains (lobes) of the sensor, governed by lobe-specific Hill kinetics:

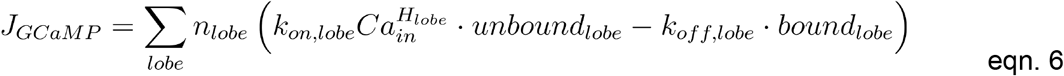

with *k*_*on,lobe*_ and *k*_*off,lobe*_, *H*_*lobe*_, and lobe specific “bound” and “unbound” terms as in **equation 1**. The leading *n*_*lobe*_ multiplier ensures strict mass conservation by accounting for the number of calcium ions bound or released per lobe-binding event, formally set as 2 for this work (though see, e.g., ^7,51^ for works that consider calmodulin domains with lobe-specific alterations to number of bound calcium).

*J*_*Buffer*_ represents the sum of the kinetic action of all buffers present in a given simulation where each buffer follows the parameters presented in **Table 2** and with form as given in their respective source references. We use the standard mass-action model to describe the calcium binding of ATP and the endogenous buffer as:

**Table 2.**
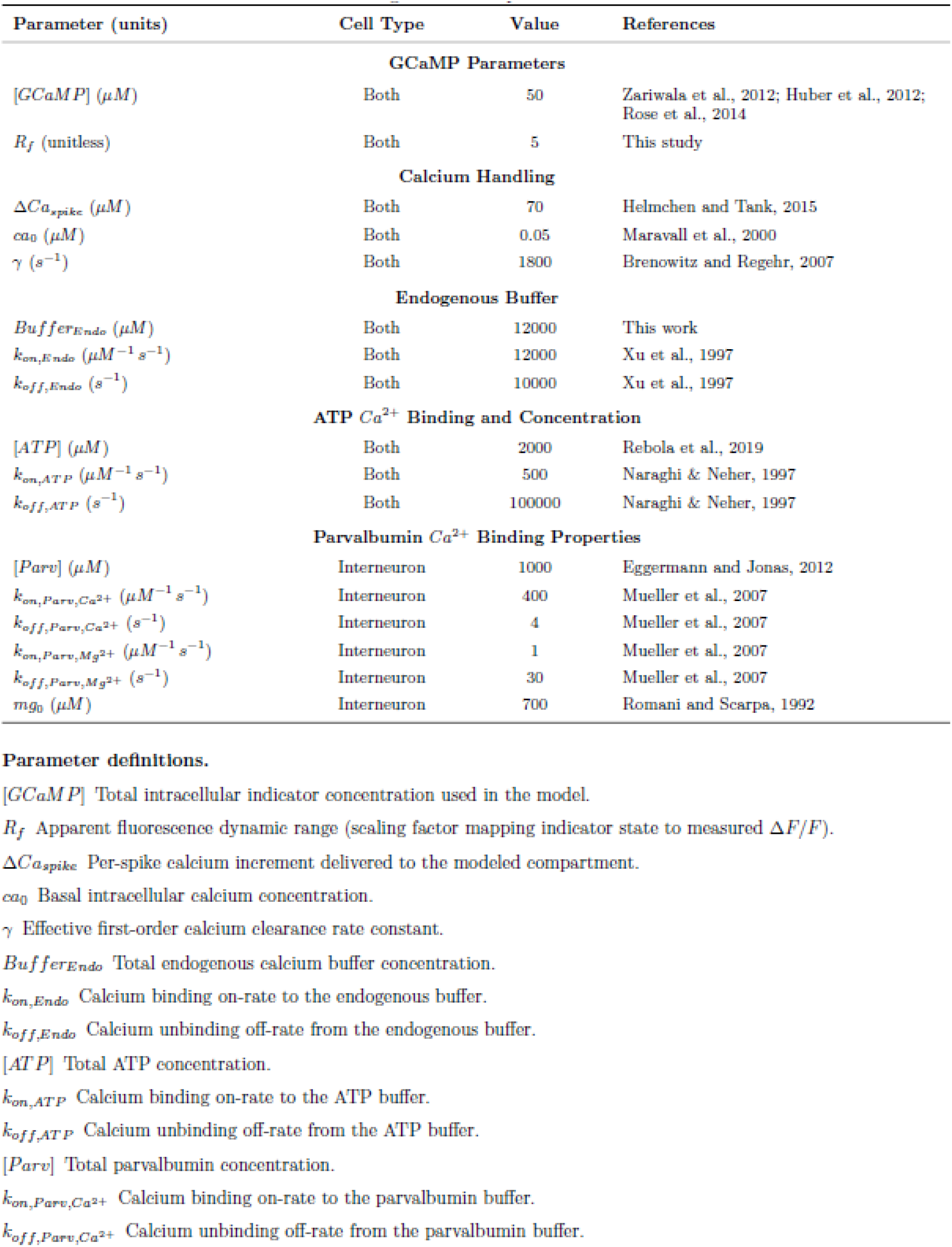
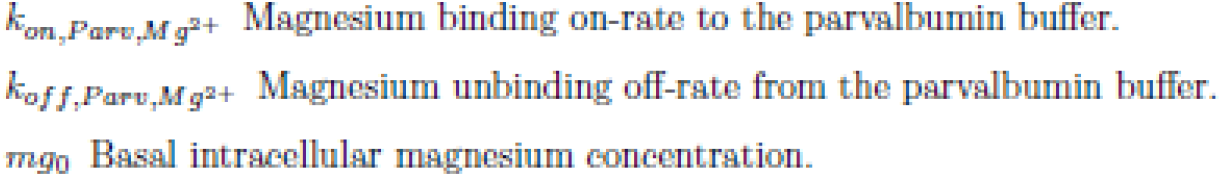
Single compartment cell model parameters.

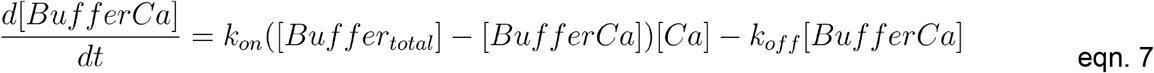

where [*Buffer*_*total*_] represents either the total concentration of endogenous buffer, [*Buffer*_*endo*_], or ATP, [*ATP*], while [*BufferCa*] represents the concentration of the buffer species bound to calcium.

To model the effects of interneuron-specific buffers, we included parvalbumin in a subset of simulations (**Supplementary Figure 8**). To model parvalbumin buffering, we follow a previous model of competitive binding between Mg2+ and Ca2+ to a shared free parvalbumin pool ^52^. The model tracked Mg-bound and Ca-bound parvalbumin states, [*ParvMg*] and [*ParvCa*], respectively. Free parvalbumin is computed from the conservation of total parvalbumin concentration, [*Parv*] as:

[*Parv*_*free*_] = [*Parv*] − [*ParvCa*] − [*ParvMg*]. This model also uses the standard mass-action model for Ca2+ and Mg2+ binding to parvalbumin.

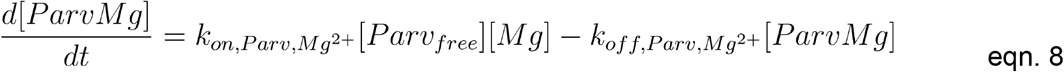

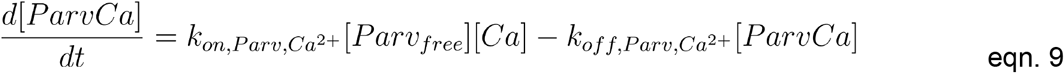

Initial concentrations of parvalbumin bound states were calculated using resting intracellular Ca2+ and Mg2+ concentrations to achieve equilibrium binding as was done in ^37^.

#### Cell Model

To simulate intracellular calcium transients (e.g., after action potentials) with minimal computational cost, we used a linearized buffer approach. The endogenous buffer capacity,, was linearized according to previous studies ^53^. Specifically, each small increment of free calcium Ca_spike_ was instantly equilibrated with endogenous buffer:

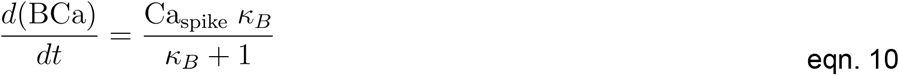

where BCa is the portion of the endogenous buffer bound to calcium, a ratio of approximately 40 for our standard granule cell model. This linearization preserves the overall calcium balance while allowing faster numerical integration of high buffer concentration simulations. It is valid under the assumptions that resting calcium (*c*_0_) is << than the affinity of the buffer for calcium (i.e., *K*_*d*_ = *k*_*offB*_ / *k*_*onB*_) and a strict separation of time scales between the endogenous buffer and explicitly modeled kinetic processes (e.g., GCaMP binding) ^54^.

The temporal evolution of intracellular free calcium *Ca*_*in*_ is then governed by the following differential equation adapted from ^37^ to include an internal calcium reservoir with influx and efflux governed by Michaelis-Menten-like dynamics :

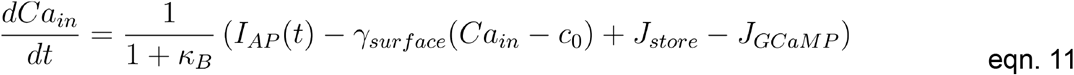

Where, *κ*_*B*_ represents the endogenous buffering capacity calculated as *κ*_*B*_ = *B*_*tot*_ / *K*_*d*_, and *I*_*AP*_ (*t*) is the calcium influx function (modeled as a delta spike).

The dynamics of the internal store (*Ca*_*store*_) contribution *J*_*store*_ are modeled as:

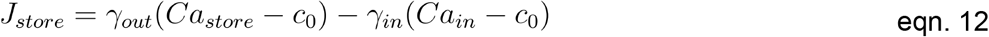

where *γ*_*in*_ is the rate constant for calcium uptake into the stores, and *γ*_*out*_ is the rate constant for calcium release from the stores. Both fluxes are driven by the difference between the respective compartment concentration and the resting concentration *c*_0_. Both the Explicit Buffer and Cell models were integrated over time using the forward Euler approach.

#### GCaMP model selection

To identify a robust set of GCaMP parameters, we employed a multi-stage optimization procedure that jointly constrained the GCaMP and cell model using both stop-flow fluorimetry and imaging datasets. We first performed gradient descent optimization initialized from a mesh of random starting parameters to generate a library of approximately 1,000 candidate parameter sets that adequately described the stopped-flow data. These candidate GCaMP models were then fixed and fit using the cell model to either our *ex vivo* or published *in vivo* recordings with known spike times as described in the text to ensure biological plausibility. Fit parameters were clustered by k-means with number of clusters set as 14 (*kmeans*, Matlab 2023b) as a way to highlight how different solutions to stopped-flow perform in cell-based contexts (**Supplementary Figure 5**). The final GCaMP model was selected by minimizing a joint objective function, defined as the sum of the z-scored L2 norms (Euclidean distance) of the fit residuals from both the target stopped-flow and imaging datasets. A collection of all GCaMP parameterizations can be found in **Table 3**.

**Table 3.**
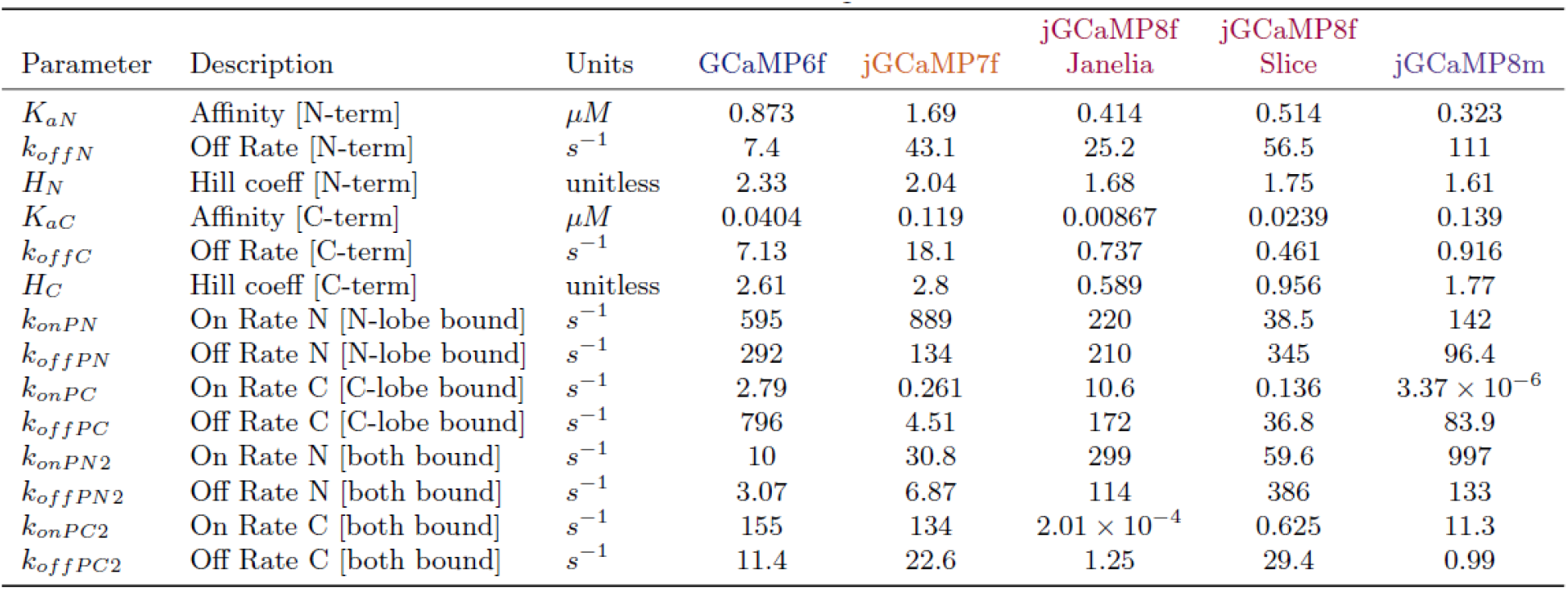
GCaMP model parameter sets.

### Bayesian and machine-learning spike inference algorithms

#### Biophys_SMC_

In order to infer spike times from fluorescence time series using the biophysical model of jGCaMP8f described in the main text we adopted the Bayesian framework developed by Diana et al, 2026 based on sequential Monte Carlo methods (particle Gibbs with ancestor sampling, PGAS; ^55^ which enabled us to estimate simultaneously the posterior distributions of spike times and the parameters of the cell model, including the GCaMP concentration, calcium extrusion rates, peak amplitude of the calcium flux due to single action potentials and the dynamic range of the probe.

Throughout this work, priors for these parameters were set as the midpoint of biologically reasonable values. (e.g. because the GCaMP concentration was allowed to vary between 1 and 120 μM, the prior was set as 60 μM). Parameter prior values and sources can be found in **Table 4**. Different priors governing dynamic range were used for jGCaMP8f versus jGCaMP8m to reflect reported differences in per-spike SNR ^5^. Specific values used in the prior set can be inspected at the C-SPIKES repository. Single point estimates of model parameters and spike times were obtained by using maximum-a-posteriori estimates. Comparisons of Linear_SMC_ were made using the PGBAR algorithm as implemented in our previous work (^26^, https://github.com/giovannidiana/pgbar) with the baseline drift parameter set to an infinitesimally small value for the PGBAR and Biophys_SMC_ algorithms (referenced throughout the text as “Linear_SMC_”, see **Supplementary Figure 7**).

**Table 4.** Cell parameter prior means and approximate ranges.

| Parameter (units) | Fixed/Free | Mean | Approximate range | References |
| --- | --- | --- | --- | --- |
| [GCaMP] ($\mu M$) | Free | 60 | 1-150 | Zariwala et al., 2012; Huber et al., 2012; Rose et al., 2014 |
| $\gamma$ ($s^{-1}$) | Free | 750 | 200-2500 | Brenowitz and Regehr, 2007 |
| $\Delta Ca_{spike}$ ($\mu M$) | Free | 15<br>5 (jGCaMP8f) | 1-100<br>3-7 (jGCaMP8f) | Helmchen and Tank, 2015 |
| $R_f$ (unitless) | Free | 10 (jGCaMP8m) | 8-12 (jGCaMP8m) | This study |
| $\gamma_{in}$ ($s^{-1}$) | Free | 5 | 0.5-50 | This study |
| $\gamma_{out}$ ($s^{-1}$) | Free | 5 | 0.5-50 | This study |
| [Buffer<sub>Endo</sub>] ($\mu M$) | Fixed | 4000 | | Rebola et al., 2019 |
| $k_{onEndo}$ ($\mu M^{-1} s^{-1}$) | Fixed | 1e2 | | Rebola et al., 2019 |
| $k_{offEndo}$ ($s^{-1}$) | Fixed | 1e4 | | Rebola et al., 2019 |

#### MLspike parameters

We compared the performance of our method with MLSpike ^15^, which employs a different model for the calcium indicator dynamics and baseline fluorescence. Spike inference in MLspike is carried out by discretizing all continuous dynamical variables in the model and applying the Viterbi algorithm to extract maximum *a-posteriori* estimates of spike times. To make a fair comparison between MLspike performance and other inference methods, we performed a sweep over tau (decay) and a (amplitude) parameters while setting *pnonlin* parameters (referenced as p2 and p3 in the original publication) to [1,0] (imposing a quadratic nonlinearity), the baseline drift was set to 0.045, the noise level was set as 0.08, and maxspike per time bin is set to 1 for the 121 Hz data processed here. MLspike outputs from the sweep were chosen on the basis of least squared error between the generative trace and the fluorescence data. Further details of the implementation can be found at our fork of the original MLspike repository (https://github.com/gerardjb/MLspike_port).

#### Supervised method comparators

We compared our method with state-of-the-art supervised machine-learning models: CASCADE ^11^ and ENS^2 12^. The two approaches use different network architectures. CASCADE returns the spike probability at each time point by taking as input a local time window by using a trained convolutional network while ENS^2^ uses the U-net applied on fixed-length segments of the original calcium trace.

In this work, we trained a CASCADE model using either the full curated dataset for the corresponding cell type from Rupprecht and colleagues (https://github.com/HelmchenLabSoftware/Cascade) or the jGCaMP8f dataset alone. For ENS^2^ comparisons, we used the published models for each cell type (https://github.com/TinLab/ENS2). Both frameworks have been updated and harmonized with the architecture of the software components included in this work’s repository. Further details of the configurations can be found in the curated collection of models maintained at the C-SPIKES repository.

#### Biophys_ML_

BiophysML is a two-stage procedure that uses Biophys_SMC_ for dataset-specific calibration, after which spike inference is performed by a supervised neural network trained on synthetic data generated from the calibrated biophysical model.

For each target dataset (indicator/cell type), we selected a representative segment of unlabeled fluorescence (typically ∼36 s; requiring ∼5 minutes to compute on an A100 GPU) and ran Biophys_SMC_ with fixed indicator kinetics (Table 2) to infer the posterior distribution over cell-specific parameters (including effective indicator concentration, calcium extrusion, per-spike calcium influx amplitude, dynamic range, and inter-compartmental rate constants). We then used the posterior mean as the calibrated parameterization for synthetic data generation. This was the only SMC computation required for Biophys_ML_.

Using the calibrated cell-parameterization, we generated paired synthetic datasets by sampling spike trains spanning the range of firing statistics relevant to the target data, forward-simulating fluorescence traces through the coupled cell/indicator model, and adding measurement noise and baseline drift to match the target dataset statistics (as in **Figure 4A**). This produces arbitrarily large, labeled fluorescence/spike pairs without requiring experimental ground truth. We then trained supervised decoders using established architectures (CASCADE and ENS^2 11,12^ on the synthetic fluorescence/spike pairs.

The key practical advantage is that Biophys_ML_ replaces long-run Monte Carlo inference with amortized neural-network inference; only the short SMC calibration step is required per indicator/cell-type while subsequent decoding runs at standard deep-learning throughput. For inference on standard datasets, we provide a curated list of pre-trained models at the C-SPIKES repository.

### Analysis of spike inference performance

#### *In vivo* data used in this study

The ground truth *in vivo* data was collected at Janelia in support of the characterization of the jGCaMP8 series of sensors ^5^. Specifically, the jGCaMP8f dataset was extracted from the Dandi dataset created by Márton Rózsa (https://dandiarchive.org/dandiset/000168). Amongst those data, we processed 154 16-second spike/fluorescence paired epochs in 37 excitatory neurons and 3 inhibitory neurons, containing 8,161 action potentials, with a median of 34 (IQR, 16-69) spikes per epoch. The jGCaMP8m dataset was extracted from the CASCADE repository (https://github.com/HelmchenLabSoftware/Cascade), with 194 ∼16 second spike/fluorescence paired epochs in 42 excitatory neurons, containing 4,575 ground truth action potentials, with a median of 16 (IQR, 8-30) spikes per epoch.

#### Analysis of continuous spike probability estimates

To evaluate the fidelity of the continuous firing rate predictions generated by the inference models (prior to discretization), we compared the inferred probability traces against ground truth electrophysiology. Ground truth spike times were converted into a continuous spike density function by convolving them with a Gaussian kernel.

We then calculated the Pearson correlation coefficient between the inferred trace and the smoothed ground truth. To characterize the temporal precision of the inference, this analysis was performed across a range of Gaussian filter widths as described in the main text. This sweep allowed us to determine the optimal temporal resolution for each method and to assess how performance degraded as the requirement for temporal precision was increased.

#### Analysis of discrete action potential imputation

Continuous prediction traces were converted to discrete imputed spike times using the approach indicated by each published method. For Biophys_SMC_, spike predictions were calculated using the maximum a-posteriori of the posterior distribution of the spike state over time. For Biophys_ML_, spike predictions were calculated by optimally fitting a set of Gaussian kernels with prior width and height determined by approximating the smoothing filter applied to ground truth for model training as in ^11^.

To reduce double-counting of imputed spikes for a given ground truth spike, we employed the linear assignment algorithm to identify unique imputed/ground truth pairs. We used the matlab function *matchpairs* at a cost of unassignment as specified in the main text to constrain the time interval between such pairs as specified in the main text. True positives were then defined as imputed spikes with unique ground truth matches, with unmatched ground truth spikes giving the false negatives, and unmatched imputed spikes being false positives. Total true positives, false positives, and ground-truth spikes within epochs for each cell to form the distributions for F-score, precision, and recall/specificity.

For each epoch of ground-truth recording, the bias in each imputed spike time was calculated as the interval to the nearest ground-truth action potential while uncertainty was defined as the absolute value of this quantity. The measures were taken to either the nearest ground truth spike using a nearest neighbors search (unmatched) or to the nearest matched spike.

### Statistics and reproducibility

Box and whisker plots comprise a box indicating the median and 25-75th percentile range and whiskers indicating the shorter of 1.5 times the interquartile range or the extreme data point. Descriptive statistics were calculated using median-based methods, and the estimated standard deviation was defined as the median absolute deviation divided by 0.6745.

To account for the hierarchical structure of our data (multiple recordings from the same animal), we fit a linear mixed-effects model to the Pearson correlation between each method’s inference output and ground truth using the *fitlme* function as implemented in MATLAB 2023b. Correlations were Fisher Z-transformed to satisfy normality and homoscedasticity assumptions of the linear mixed-effects model as correlation values skewed left. The model included method and either sampling frequency or filter width (as indicated in the main text) as fixed effects, and animalId/epoch as a random intercept. Statistical significance of fixed effects was assessed using the Wald F-Test to test for main effect. Post-hoc pairwise comparisons using a linear contrast with Bonferroni correction were then conducted as described in the main text.

## Acknowledgements

The authors thank Marissa Applegate, Kiri Couchman, and Minglu Wang for collaboration and technical assistance in early stages of this project, Yan Zhang and Loren Looger for discussion and for providing a jGCaMP8f construct prior to publication, and Jinho Park for careful reading and discussion of the manuscript. S.W. was supported by NIH R01 NS045193 and U19 NS104648. S.W. and D.D. were supported by NIH R21 EY026434. G.J.B. was supported by the Warren Alpert Foundation and the Brain & Behavior Research Foundation. The laboratory of D.D. was also supported by the Fondation pour la Recherche Médicale (FRM EQU202003010555), Fondation pour l’Audition (FPA-RD-2018-8), and the Agence Nationale de la Recherche (ANR-17-CE16-0019, and ANR-18-CE16-0018, ANR-19-CE16 0019-02, ANR-21-CE16-0036-01).

## Code availability

Code is available on github at https://github.com/gerardjb/C-SPIKES. This includes implementations of the Biophys_SMC_ and Biophys_ML_ methods, curated Biophys_ML_ models, and tools for comparing and visualizing performance of the biophysical methods with other comparator methods.

## References

1. Tian, L. et al. Imaging neural activity in worms, flies and mice with improved GCaMP calcium indicators. Nat. Methods 6, 875 (2009).

2. Akerboom, J. et al. Optimization of a GCaMP calcium indicator for neural activity imaging. J. Neurosci. Off. J. Soc. Neurosci. 32, 13819–13840 (2012).

3. Chen, T.-W. et al. Ultrasensitive fluorescent proteins for imaging neuronal activity. Nature 499, 295–300 (2013).

4. Dana, H. et al. High-performance calcium sensors for imaging activity in neuronal populations and microcompartments. Nat. Methods 16, 649–657 (2019).

5. Zhang, Y. et al. Fast and sensitive GCaMP calcium indicators for imaging neural populations. Nature 615, 884–891 (2023).

6. Helmchen, F. & Tank, D. W. A single-compartment model of calcium dynamics in nerve terminals and dendrites. Cold Spring Harb. Protoc. 2015, 155–167 (2015).

7. Sun, X. R. et al. Fast GCaMPs for improved tracking of neuronal activity. Nat. Commun. 4, 2170 (2013).

8. Xiao, D. et al. Mapping cortical mesoscopic networks of single spiking cortical or sub-cortical neurons. eLife 6, e19976 (2017).

9. Li, P. et al. Measuring Sharp Waves and Oscillatory Population Activity With the Genetically Encoded Calcium Indicator GCaMP6f. Front. Cell. Neurosci. 13, (2019).

10. Wei, Z. et al. A comparison of neuronal population dynamics measured with calcium imaging and electrophysiology. PLOS Comput. Biol. 16, e1008198 (2020).

11. Rupprecht, P. et al. A database and deep learning toolbox for noise-optimized, generalized spike inference from calcium imaging. Nat. Neurosci. 24, 1324–1337 (2021).

12. Zhou, Z. et al. Effective and efficient neural networks for spike inference from in vivo calcium imaging. Cell Rep. Methods 3, 100462 (2023).

13. Vogelstein, J. T. et al. Fast Nonnegative Deconvolution for Spike Train Inference From Population Calcium Imaging. J. Neurophysiol. 104, 3691–3704 (2010).

14. Pnevmatikakis, E. A., Merel, J., Pakman, A. & Paninski, L. Bayesian spike inference from calcium imaging data. Preprint at 10.48550/arXiv.1311.6864 (2013).

15. Deneux, T. et al. Accurate spike estimation from noisy calcium signals for ultrafast three-dimensional imaging of large neuronal populations in vivo. Nat. Commun. 7, 12190 (2016).

16. Friedrich, J., Zhou, P. & Paninski, L. Fast online deconvolution of calcium imaging data. PLOS Comput. Biol. 13, e1005423 (2017).

17. Sebastian, J. et al. GDspike: An accurate spike estimation algorithm from noisy calcium fluorescence signals. 2017 IEEE Int. Conf. Acoust. Speech Signal Process. ICASSP 1043–1047 (2017) doi:10.1109/ICASSP.2017.7952315.

18. Greenberg, D. S. et al. Accurate action potential inference from a calcium sensor protein through biophysical modeling. 479055 Preprint at 10.1101/479055 (2018).

19. Pachitariu, M., Stringer, C. & Harris, K. D. Robustness of Spike Deconvolution for Neuronal Calcium Imaging. J. Neurosci. 38, 7976–7985 (2018).

20. Theis, L. et al. Benchmarking Spike Rate Inference in Population Calcium Imaging. Neuron 90, 471–482 (2016).

21. Regehr, W. G. & Atluri, P. P. Calcium transients in cerebellar granule cell presynaptic terminals. Biophys. J. 68, 2156–2170 (1995).

22. Sabatini, B. L. & Regehr, W. G. Optical measurement of presynaptic calcium currents. Biophys. J. 74, 1549–1563 (1998).

23. Rose, T., Goltstein, P. M., Portugues, R. & Griesbeck, O. Putting a finishing touch on GECIs. Front. Mol. Neurosci. 7, (2014).

24. Seroussi, I. & Zeitouni, O. Lower Bounds on the Generalization Error of Nonlinear Learning Models. Preprint at 10.48550/arXiv.2103.14723 (2022).

25. Sariyildiz, M. B., Kalantidis, Y., Alahari, K. & Larlus, D. No Reason for No Supervision: Improved Generalization in Supervised Models. Preprint at 10.48550/arXiv.2206.15369 (2023).

26. Diana, G., Sermet, B. S., Broussard, G. J., Wang, S. S.-H. & DiGregorio, D. A. High frequency spike inference with particle Gibbs sampling. eLife 13, (2026).

27. Éltes, T., Szoboszlay, M., Kerti-Szigeti, K. & Nusser, Z. Improved spike inference accuracy by estimating the peak amplitude of unitary [Ca2+] transients in weakly GCaMP6f-expressing hippocampal pyramidal cells. J. Physiol. 597, 2925–2947 (2019).

28. Huang, L. et al. Relationship between simultaneously recorded spiking activity and fluorescence signal in GCaMP6 transgenic mice. eLife 10, e51675 (2021).

29. Schoenfeld, G., Carta, S., Rupprecht, P., Ayaz, A. & Helmchen, F. In Vivo Calcium Imaging of CA3 Pyramidal Neuron Populations in Adult Mouse Hippocampus. eNeuro 8, (2021).

30. Brenowitz, S. D. & Regehr, W. G. Reliability and Heterogeneity of Calcium Signaling at Single Presynaptic Boutons of Cerebellar Granule Cells. J. Neurosci. 27, 7888–7898 (2007).

31. Badura, A., Sun, X. R., Giovannucci, A., Lynch, L. A. & Wang, S. S. H. Fast calcium sensor proteins for monitoring neural activity. Neurophotonics 1, 025008 (2014).

32. Linse, S., Helmersson, A. & Forsén, S. Calcium binding to calmodulin and its globular domains. J. Biol. Chem. 266, 8050–8054 (1991).

33. Faas, G. C., Raghavachari, S., Lisman, J. E. & Mody, I. Calmodulin as a direct detector of Ca2+ signals. Nat. Neurosci. 14, 301–304 (2011).

34. Johnson, J. D., Snyder, C., Walsh, M. & Flynn, M. Effects of Myosin Light Chain Kinase and Peptides on Ca2+ Exchange with the N- and C-terminal Ca2+ Binding Sites of Calmodulin (*). J. Biol. Chem. 271, 761–767 (1996).

35. Brown, S. E., Martin, S. R. & Bayley, P. M. Kinetic Control of the Dissociation Pathway of Calmodulin-Peptide Complexes*. J. Biol. Chem. 272, 3389–3397 (1997).

36. Lai, M., Brun, D., Edelstein, S. J. & Novère, N. L. Modulation of Calmodulin Lobes by Different Targets: An Allosteric Model with Hemiconcerted Conformational Transitions. PLOS Comput. Biol. 11, e1004063 (2015).

37. Rebola, N. et al. Distinct Nanoscale Calcium Channel and Synaptic Vesicle Topographies Contribute to the Diversity of Synaptic Function. Neuron 104, 693-710.e9 (2019).

38. Speiser, A. et al. Fast amortized inference of neural activity from calcium imaging data with variational autoencoders. in Advances in Neural Information Processing Systems vol. 30 (Curran Associates, Inc., 2017).

39. Inoue, M. et al. Rational design of a high-affinity, fast, red calcium indicator R-CaMP2. Nat. Methods 12, 64–70 (2015).

40. Inoue, M. et al. Rational Engineering of XCaMPs, a Multicolor GECI Suite for In Vivo Imaging of Complex Brain Circuit Dynamics. Cell 177, 1346-1360.e24 (2019).

41. Dana, H. et al. Sensitive red protein calcium indicators for imaging neural activity. eLife 5, e12727 (2016).

42. Deo, C. et al. The HaloTag as a general scaffold for far-red tunable chemigenetic indicators. Nat. Chem. Biol. 17, 718–723 (2021).

43. Farrants, H. et al. A modular chemigenetic calcium indicator for multiplexed in vivo functional imaging. Nat. Methods 21, 1916–1925 (2024).

44. Evans, S. W. et al. A positively tuned voltage indicator for extended electrical recordings in the brain. Nat. Methods 20, 1104–1113 (2023).

45. Platisa, J. et al. High-speed low-light in vivo two-photon voltage imaging of large neuronal populations. Nat. Methods 20, 1095 (2023).

46. Patriarchi, T. et al. Ultrafast neuronal imaging of dopamine dynamics with designed genetically encoded sensors. Science 360, (2018).

47. Sun, F. et al. A Genetically Encoded Fluorescent Sensor Enables Rapid and Specific Detection of Dopamine in Flies, Fish, and Mice. Cell 174, 481-496.e19 (2018).

48. Tran, V., Park, M. C. H. & Stricker, C. An improved measurement of the Ca2+-binding affinity of fluorescent Ca2+ indicators. Cell Calcium 71, 86–94 (2018).

49. Ward, W. W. Biochemical and Physical Properties of Green Fluorescent Protein. in Green Fluorescent Protein 39–65 (John Wiley & Sons, Ltd, 2005). doi:10.1002/0471739499.ch3.

50. Barnett, L. M., Hughes, T. E. & Drobizhev, M. Deciphering the molecular mechanism responsible for GCaMP6m’s Ca2+-dependent change in fluorescence. PLOS ONE 12, e0170934 (2017).

51. Helassa, N. et al. Fast-Response Calmodulin-Based Fluorescent Indicators Reveal Rapid Intracellular Calcium Dynamics. Sci. Rep. 5, 15978 (2015).

52. Müller, M., Felmy, F., Schwaller, B. & Schneggenburger, R. Parvalbumin is a mobile presynaptic Ca2+ buffer in the calyx of Held that accelerates the decay of Ca2+ and short-term facilitation. J. Neurosci. Off. J. Soc. Neurosci. 27, 2261–2271 (2007).

53. Neher, E. & Augustine, G. J. Calcium gradients and buffers in bovine chromaffin cells. J. Physiol. 450, 273–301 (1992).

54. Wagner, J. & Keizer, J. Effects of rapid buffers on Ca2+ diffusion and Ca2+ oscillations. Biophys. J. 67, 447–456 (1994).

55. Lindsten, F., Jordan, M. I. & Schön, T. B. Particle Gibbs with Ancestor Sampling. J. Mach. Learn. Res. 15, 2145–2184 (2014).

